# A gene regulatory network for neural induction

**DOI:** 10.1101/2021.04.16.440164

**Authors:** Katherine E. Trevers, Hui-Chun Lu, Youwen Yang, Alexandre Thiery, Anna C. Strobl, Božena Pálinkášová, Nidia M. M. de Oliveira, Irene M. de Almeida, Mohsin A. F. Khan, Natalia Moncaut, Nicholas M. Luscombe, Leslie Dale, Andrea Streit, Claudio D. Stern

## Abstract

During early vertebrate development, signals from a special region of the embryo, the organizer, can re-direct the fate of non-neural ectoderm cells to form a complete, patterned nervous system. This is called neural induction and has generally been imagined as a single signalling event, causing a switch of fate. Here we undertake a comprehensive analysis, in very fine time-course, of the events following exposure of ectoderm to the organizer. Using transcriptomics and epigenomics we generate a Gene Regulatory Network comprising 175 transcriptional regulators and 5,614 predicted interactions between them, with fine temporal dynamics from initial exposure to the signals to expression of mature neural plate markers. Using in situ hybridization, single-cell RNA-sequencing and reporter assays we show that neural induction by a grafted organizer closely resembles normal neural plate development. The study is accompanied by a comprehensive resource including information about conservation of the predicted enhancers in different vertebrate systems.

## Introduction

One of the most influential studies in developmental biology was the discovery, 100 years ago, that a small region of the vertebrate embryo, named the “organizer”, can induce ectodermal cells that do not normally contribute to the neural plate to form a complete, patterned nervous system (Spemann, 1921; Spemann & Mangold, 1924). In amphibians, where these experiments were initially conducted, the “organizer” resides in the dorsal lip of the blastopore. A few years later, Waddington demonstrated that an equivalent region exists in birds (duck and chick) and mammals (rabbit) (Waddington, 1933, 1934, 1936, 1937; Waddington & Schmidt, 1933). In these groups, an equivalent organizer is located at the tip of the primitive streak, a structure known as Hensen’s node (Hensen, 1876). This interaction between the organizer and the responding ectoderm, which causes the latter to acquire neural plate identity, has been termed “neural induction”(Gallera, 1971b; Gurdon, 1987; Nieuwkoop et al., 1952; Saxen, 1980; Spemann & Mangold, 1924; Stern, 2005; Storey, Crossley, De Robertis, Norris, & Stern, 1992).

Neural induction has often been imagined as a single event, “switching” the fate of the responding tissue from non-neural to neural. But it is clear from timed transplantation experiments that the responding ectoderm requires several hours’ (about 12) exposure to the organizer (Gallera, 1971b; Gallera & Ivanov, 1964; Streit et al., 1998) to acquire its new identity in a stable way (“commitment”). Recent work has also revealed that the expression of many genes changes over time after grafting an organizer, suggesting that the process has considerable complexity (Albazerchi & Stern, 2007; Gibson, Robinson, Streit, Sheng, & Stern, 2011; Costis Papanayotou et al., 2013; C. Papanayotou et al., 2008; Pinho et al., 2011; Sheng, dos Reis, & Stern, 2003; Stern, 2005; Streit, Berliner, Papanayotou, Sirulnik, & Stern, 2000; Streit et al., 1998; Streit et al., 1997; Streit & Stern, 1999; Trevers et al., 2018). What happens during this period? Is it possible to define distinct steps, and perhaps the molecular events that represent “induction” and “commitment”? Surprisingly, given the long time since the discovery of neural induction, these questions have hardly been addressed. To begin to answer them requires a comprehensive approach to identify and model the key interactions between genes and transcriptional regulators inside the responding cells that accompany their responses to signals from the organizer over time.

Grafting an organizer to a site that does not normally contribute to neural tissue but is competent to do so provides the opportunity to study the progression of neural induction relative to “time-0”, the moment when the organizer is first presented to the tissue. This allows for the separation of neural inductive events from other processes that occur during normal neural plate development (for example mesendoderm formation and patterning), because sites that are remote enough from the normal neural plate and are competent to respond to an organizer graft can generate a complete neural tube without induction of mesoderm (Dias & Schoenwolf, 1990; Hornbruch, Summerbell, & Wolpert, 1979; Storey et al., 1992).

Here we have taken advantage of these properties, together with recent major technological advances in transcriptomics and epigenetic analysis, to undertake a comprehensive dissection of the molecular events that accompany neural induction in fine time-course and to generate the first detailed Gene Regulatory Network (GRN) for this process. The GRN comprises 175 transcriptional regulators and 5,614 predicted interactions between them over the course of neural induction. We then compare the spatial and temporal properties of these changes with development of the normal neural plate of the embryo using *in situ* hybridization, single cell RNA-sequencing and reporter assays to test the activity of some key gene regulatory elements. We present a comprehensive resource allowing visualization and querying of all of these interactions and regulatory elements on a genome-wide level. Together, our study offers a comprehensive view of the genetic hierarchy of neural induction and normal neural plate development over time.

## Results

### Transcriptional profiling identifies responses to neural induction in time course

In the chick embryo, a graft of Hensen’s node to an extraembryonic region of competent non-neural ectoderm, the inner area opaca (Dias & Schoenwolf, 1990; Gallera & Ivanov, 1964; Storey et al., 1992; Streit et al., 1998), induces a mature, patterned neural tube after 15h of culture (Figure 1A). To characterize the events over this period, the induction of several neural markers was assessed. *OTX2* is first expressed in the pre-primitive-streak stage epiblast at EGKXII-XIII (Albazerchi & Stern, 2007; Pinho et al., 2011) and is induced by 7/7 grafts after 3h (Figure 1B-F). *SOX2* is first expressed at HH4+/5 (Rex et al., 1997; Sheng et al., 2003; Streit et al., 1998; Streit et al., 1997) and requires 9h for induction (8/8) (Figure 1G-K). *SOX1* starts to be expressed weakly in the neural plate around HH7-8 (Stavridis, Collins, & Storey, 2010; Uchikawa et al., 2011) and is induced by 50% of grafts after 12h (Figure 1L-P). These time courses confirm that a sequence of events occurs in response to grafted nodes over 0-12h (Figure S1A), culminating in the acquisition of neural plate/tube identity (Albazerchi & Stern, 2007; Gibson et al., 2011; Costis Papanayotou et al., 2013; C. Papanayotou et al., 2008; Pinho et al., 2011; Sheng et al., 2003; Stern, 2005; Streit et al., 2000; Streit et al., 1998; Streit et al., 1997; Streit & Stern, 1999; Trevers et al., 2018).

**Figure 1.**
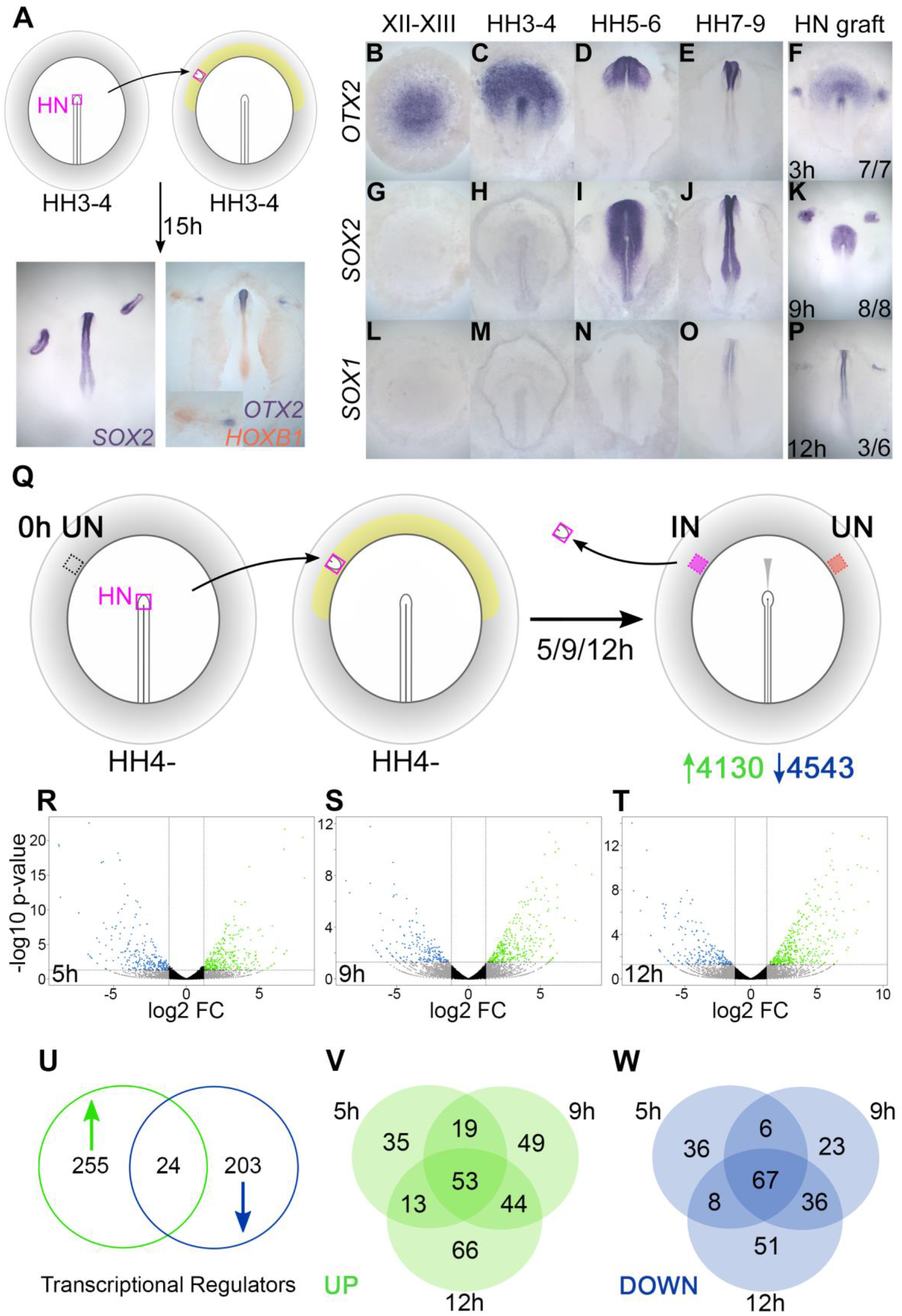
Transcriptional profiling identifies responses to neural induction in time course. (A) Hensen’s node (HN) was grafted from HH3-4 donors to the inner area opaca (yellow) of hosts at the same stage. Ectopic neural tubes expressing *SOX2*, *OTX2* and *HOXB1* are induced after 15h of culture. (B-F) Expression of neural markers compared to their time course of induction by a node graft: *OTX2*; first expressed in pre-streak epiblast (EGKXII-XIII) and induced by grafts after 3h. (G-K) *SOX2*; first expressed in the neural plate at HH5-6 and induced after 9h. (L-P) *SOX1*; first expressed in the forming neural tube at HH7-8 and induced after 12h (3/6 cases). (Q) Identifying transcriptional responses to a node graft. HN was grafted from HH4-donors to HH4-hosts. The HN graft was removed and underlying “Induced” (IN) and contralateral “Uninduced” (UN) ectoderm isolated after 5, 9 or 12h. Uninduced “0h” ectoderm from HH4-embryos was also dissected. Samples were processed by RNAseq. (R-T) Induced and corresponding uninduced tissues were compared at each time point to identify differentially expressed transcripts. Volcano plots show upregulated (green) and downregulated (blue) transcripts (> ±1.2 log_2_ fold change and <0.05 p-value). (U-W) Venn diagrams represent 482 genes encoding transcriptional regulators (U) that are upregulated (V) or downregulated (W) at different time points.

To uncover the full transcriptional responses to neural induction over this period, we performed RNAseq at three time points following Hensen’s node grafts: 5h (to identify early “pre-neural” responses), 9h (when *SOX2* expression defines neural specification) and 12h, as cells start to express *SOX1*. Hensen’s node was grafted from HH4-chick donors to a region of competent epiblast (inner area opaca) of chick hosts at the same stage. Embryos were cultured for the desired period of time, after which the graft was removed and the underlying “Induced” tissue was collected, as well as control (“Uninduced”) ectoderm from the corresponding region on the contralateral side of the same embryos (Figure 1Q). Competent area opaca at HH4-, representing a “0h” starting point for the time course, was also sampled. Transcripts were sequenced and mapped to the chicken genome. When comparing “Induced” and “Uninduced” reads at each time point, DESeq analyses identified 8673 differentially expressed transcripts based on a ±1.2 log_2_ fold change (Figure 1R-T). Of these, 4130 were upregulated (enriched in “induced” tissue) and 4543 were downregulated (depleted in “induced” tissue) relative to the “uninduced” counterpart.

To construct a GRN we focused on 482 transcription factors and chromatin modifiers that are differentially expressed within these data. Of these “transcriptional regulators”, 255 are upregulated, 203 are downregulated and 24 have more complex expression dynamics (Figures 1U and S1B). Grouping transcripts based on the timing of their response (Figure 1V-W) reveals that 120 are differentially expressed throughout (53 upregulated and 67 downregulated), while others are unique to particular time points. Therefore, the responses to a graft of the organizer comprise changes in the expression of many genes over a relatively short period, and are therefore complex and dynamic.

Due to this complexity, we chose to increase the time resolution of the analysis. NanoString nCounter was used to quantify the gene expression changes of transcriptional regulators at 6 time points (1, 3, 5, 7, 9 and 12h after the node graft) in “induced” tissue compared to “uninduced” ectoderm. By consolidating the data from RNAseq and NanoString, a set of refined expression profiles was established for 156 transcriptional regulators that are enriched and 57 that are depleted in induced tissues (Tables S1-4). These represent the core components of our GRN.

### Epigenetic changes identify chromatin elements associated with neural induction

We next sought to identify the regions of non-coding chromatin that govern responses to signals from the organizer. Histone modifications regulate gene expression by altering chromatin structure. For example, H3K27ac is associated with actively transcribed genes and their enhancers, whereas transcriptionally inactive regions are often marked by H3K27me3 (Bonn et al., 2012; Creyghton et al., 2010; Heintzman et al., 2009; Kharchenko et al., 2011; Rada-Iglesias et al., 2011; Tiwari et al., 2008; Tolhuis et al., 2011; Zentner, Tesar, & Scacheri, 2011). To detect H3K27 chromatin marks that change during neural induction, ChIPseq was performed on induced and uninduced ectoderm following 5, 9 or 12h of a node graft (Figure 2A).

**Figure 2.**
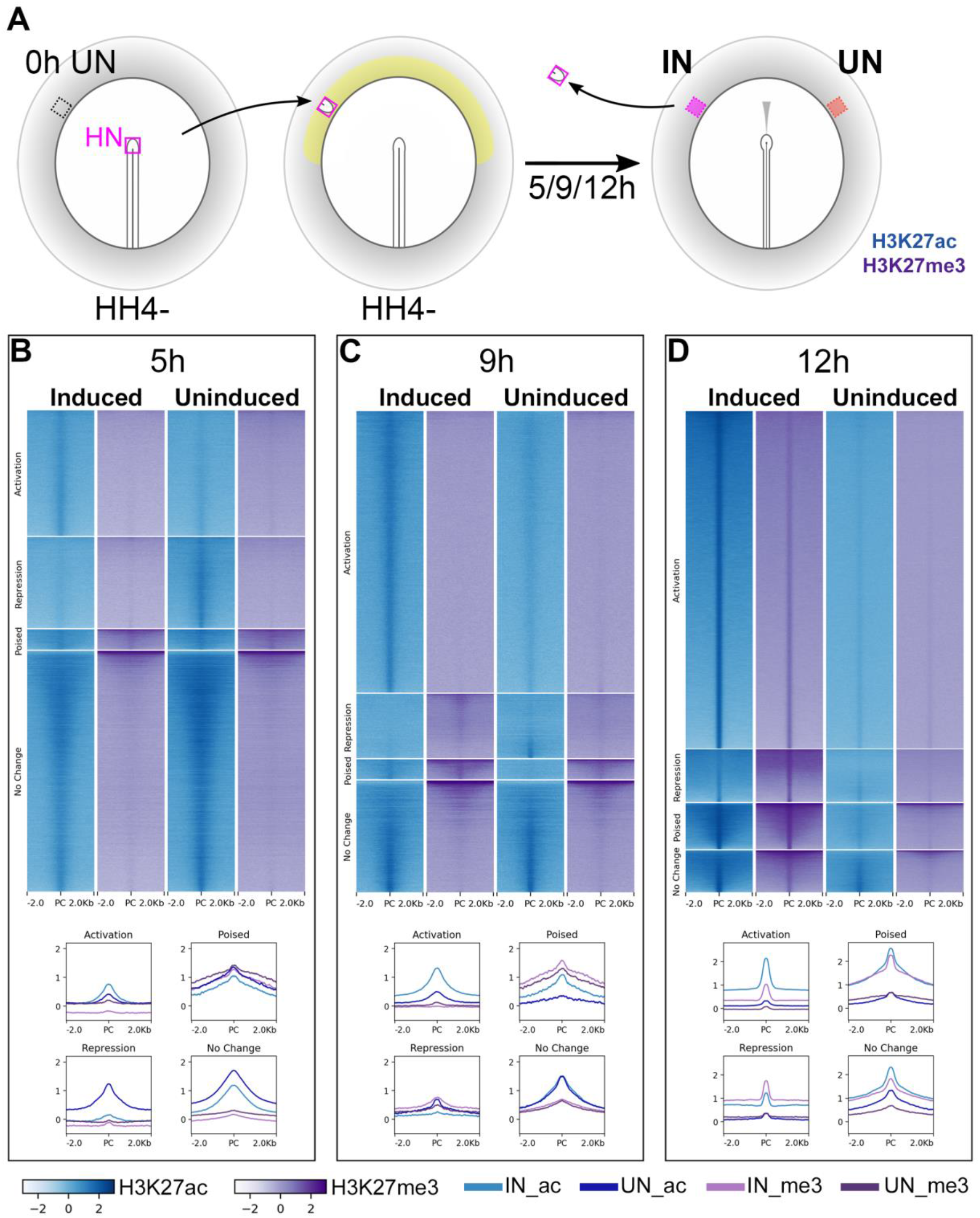
Epigenetic changes identify chromatin elements associated with neural induction. (A) Hensen’s node (at HH4-) was grafted to hosts of the same stage. Embryos were cultured for 5, 9 or 12h before HN was removed and Induced (IN) and contralateral uninduced (UN) ectoderm collected. ChIPseq was performed for H3K27ac and H3K27me3. (B-D) Putative regulatory sites were predicted according to the H3K27 profiles of IN and UN tissues at each time point (see Fig S2A). They include sites undergoing “Activation” or “Repression”, being “Poised”, or showing no change. Heat maps illustrate the enrichment of H3K27ac (blue) and H3K27me3 (purple) in IN and UN tissues within ±2.0kb from the peak centre (PC) for each annotated group. Graphs plot the distributions of H3K27ac and H3K27me3 enrichment for each group.

H3K27ac and H3K27me3 enriched regions were identified genome-wide by comparison to a genomic input sample. Chromatin sites were categorized according to their histone signatures across induced and corresponding uninduced tissues at each time point (Figure S2A). Sites that become acetylated and/or demethylated in induced tissues compared to uninduced were considered to undergo “activation” (Indices 1-3); those that become deacetylated and/or methylated undergo “repression” (Indices 4-6). Chromatin marked by both H3K27ac (activation) and H3K27me3 (repression) marks in either tissue were described as “Poised” (Indices 7-12). The remaining indices (13-16) define sites that do not change marks between the two conditions.

Chromatin sites undergoing activation are enriched for H3K27ac marks in induced tissues at each time point (Figure 2B-D). Sites of repression are more varied: those belonging to this category at 5h lose acetylation, whereas a number of sites gain methylation in induced tissues at 9h and 12h. Very few sites are “poised” at any time. Overall, as neural induction progresses, the number of sites undergoing activation increases, along with a reduction in those that do not change state.

Next, we focused on changes associated with the 213 genes encoding transcriptional regulators for which refined expression profiles were established by NanoString analysis (see above). “Constitutive” CTCF-bound sites (putative insulators) flanking these genes were obtained from chicken CTCF-ChIPseq data (Kadota et al., 2017; Khan et al., 2013). They were used to predict the boundaries within which H3K27 marks were identified by ChIPseq; these regions (“loci”) are up to 500kb in length and may contain several genes (Figure S2B). Within each locus, the H3K27ac or H3K27me3 enriched peaks were categorized as before (Figure S2A). The 213 selected loci contain a total of 6,971 sites that change in response to a node graft (Indices 1-10; Figure S2A). Our analysis identifies these as putative regulatory elements that are controlled epigenetically during the process.

### A Gene Regulatory Network for neural induction

Having identified transcriptional responses to signals from the organizer and their accompanying chromatin changes, we combined them to construct a GRN to describe the time course of these events and to illustrate predicted interactions between transcriptional regulators. The open source platform BioTapestry (Longabaugh, Davidson, & Bolouri, 2005, 2009; Paquette, Leinonen, & Longabaugh, 2016) is now used extensively to provide a visually intuitive representation of developmental networks (Arnone & Davidson, 1997; Betancur, Bronner-Fraser, & Sauka-Spengler, 2010; Davidson, 1990; Davidson, Cameron, & Ransick, 1998; Simoes-Costa & Bronner, 2015; Thiery, Buzzi, & Streit, 2020). Importantly, this software allows visualization of dynamic changes in the interactions between regulators and their target genes.

### A BioTapestry model for regulatory gene interactions

A custom computational pipeline was developed to integrate these complex time course data (Figure 3A). The 213 transcriptional regulators were selected as candidate components of the network based on their responses to a node graft in RNAseq and NanoString, and associated chromatin changes according to ChIPseq. We aimed to generate a GRN for interactions occurring during neural induction. The interactions were predicted by screening of transcription factor binding sites within the putative regulatory sites with indices 1-3 and 7 (Figure S2C-E) associated with a member of the GRN.

**Figure 3.**
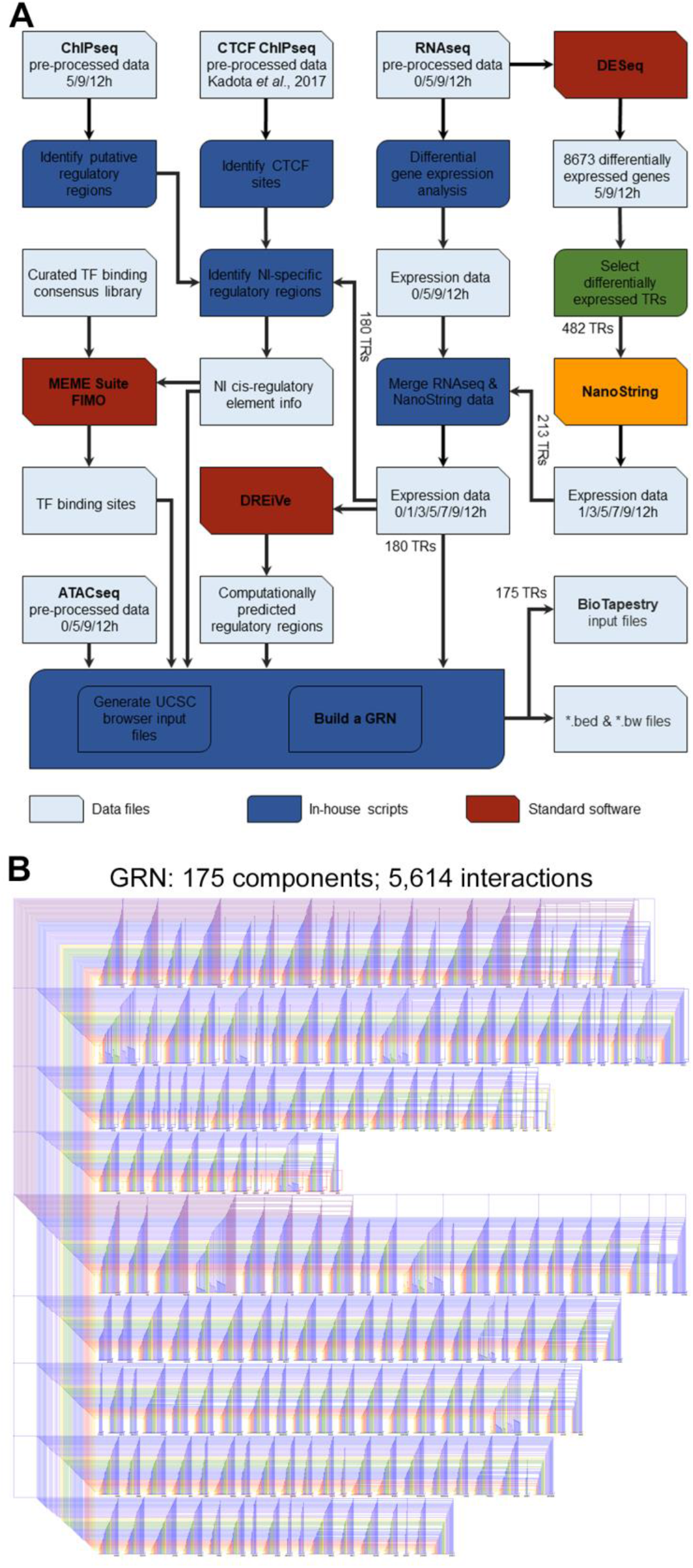
Constructing a BioTapestry model for regulatory gene interactions. (A) Computational pipeline to combine transcriptomic and epigenetic time course data, to build a GRN for neural induction. The output data are available to view in the UCSC browser. (B) A GRN comprising 5,614 predicted interactions between 175 components is visualized using BioTapestry.

Positive or negative regulations were then modelled between regulators and their putative targets by overlaying the predicted transcription factor binding profiles with consolidated expression profiles from RNAseq and NanoString using the regulatory rules shown in Figure S3A-B. At each time point, changes in expression of a putative regulator that could lead to the same change in a candidate target are modelled as positive interactions. Negative, or inhibitory, interactions are predicted when changes in expression of the regulator could cause the opposite change in a downstream target. Genes lacking both regulatory inputs and outputs with other genes in the network were excluded.

The resulting GRN, depicting interactions between 175 genes, was represented using BioTapestry (Figure 3B and Suppl File S1). Most targets are predicted to receive multiple positive and/or negative inputs, which are initiated at, and can act across, multiple time points. For example, the non-neural gene *GATA2* is predicted to be repressed by *TFAP2C* after just 1h of a node graft, followed by increasing repression via *HIF1A*, *MEIS2* and numerous other factors over the remaining time course. On the other hand, *BLIMP1* is predicted to be induced by *TFAP2C* after 1h, alongside *SNAI1*, *ETV1*, *ETV4*, *MGA* and *SOX4* (Suppl File S2). In total, 5,614 interactions are predicted to occur between these components, highlighting the intricate and highly dynamic sequence of regulatory events that are triggered by signals from the organizer.

### Incorporating individual regulatory sites and their dynamics into the network

BioTapestry networks usually represent each gene once, with multiple regulatory inputs, and each regulator as a single input into the target gene. However, it is now known that gene expression is controlled by multiple regulatory elements, each with characteristic spatial and temporal activity. In the past, such elements were identified by constructing many reporter fragments by nested deletions, whose activity was then tested *in vivo*; for example, the pioneering work of H. Kondoh’s lab in identifying many enhancers controlling *SOX2* expression (Okamoto, Uchikawa, & Kondoh, 2015; Uchikawa, Ishida, Takemoto, Kamachi, & Kondoh, 2003). Our epigenetic analysis reveals that in almost all cases there are indeed multiple putative regulatory sites associated with genes that change expression, which can be in different chromatin states (Figure 4A). By predicting the changes that occur as sites undergo epigenetic “activation” or “repression” (using Indices 1-7; Figure S3A), we can model the different contributions of these chromatin elements. Since BioTapestry lacks a notation for distinct regulatory elements of individual genes, as an example, we generated a sub-network showing 6 candidate *cis*-regulatory regions of *BRD8* at 5, 9 and 12h as separate targets, where each target is represented with the BioTapestry notation used for genes (Figure 4B-D). This reveals which specific elements contain binding sites for each regulating transcription factor. One element (site1 in Figure 4B) is initially active, however it does not receive inputs from other GRN components. Site1 is then methylated at 9h and 12h as an increasing number of other elements become active, presumably to stabilize and maintain *BRD8* expression. Although it is not practical to generate a full BioTapestry representation of the entire network including all the elements in their different states in this way, all identified elements and their predicted activity are presented in the associated UCSC Browser tracks (https://genome.ucsc.edu/s/stern_lab/Neural_Induction_2021) together with the predicted binding sites relevant to the network contained within each site; a full list is also given in Tables S5-S7.

**Figure 4.**
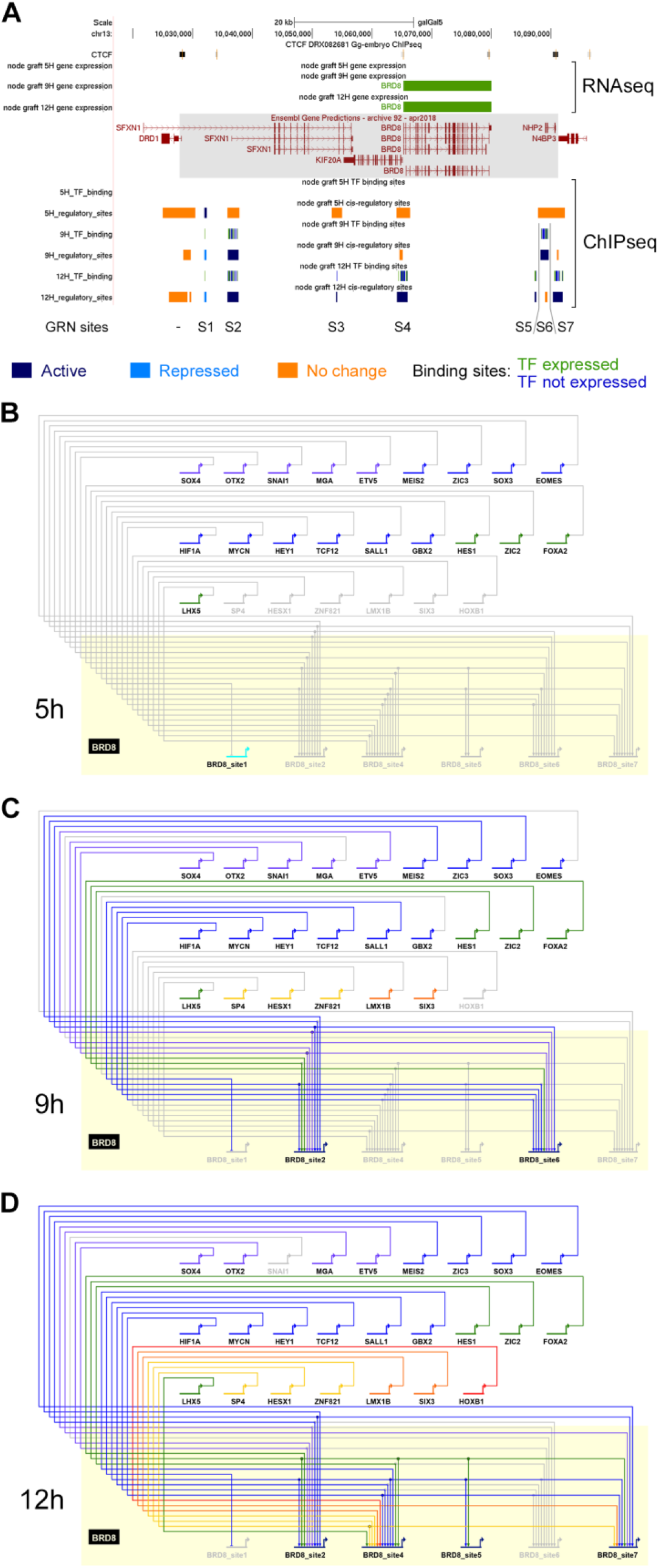
A subnetwork incorporating individual regulatory sites and their dynamics. (A) UCSC browser view of RNAseq and ChIPseq tracks associated with *BRD8*. *BRD8* is upregulated after 9h and 12h of neural induction (green bars). The *BRD8* regulatory locus (grey box; chr13:10028086-10091079) is defined by flanking CTCF-bound sites. Seven putative regulatory sites (S1-7) within this domain were predicted based on the ChIPseq H3K27 profiles. This includes sites that undergo activation (indices 1-3, 7, coloured in dark blue), repression (indices 4-6, coloured in cyan) or show no change (coloured in orange). Transcription factor binding sites by network components are shown; green for components that are expressed at the same time point and blue for those that are not. A BioTapestry subnetwork was generated from these predicted binding sites and expression profiles. Site 3, active at 12h, is not shown in the subnetwork as there is no TF that is expressed at 12h and predicted to bind to it. (B) *BRD8* regulation during neural induction: *BRD8* is initially not expressed; site 1 is active but is not predicted to be bound by other GRN components. (C) *BRD8* is upregulated after 9h; sites 2 and 6 undergo activation and could be bound by various transcription factors that are also expressed. Site 1 undergoes repression. (D) *BRD8* expression is maintained at 12h; regulators potentially bind to sites 2, 4, 5 and 7. Site 6 is no longer predicted to be active.

### Neural induction by a graft of Hensen’s node recapitulates normal neural plate development

To what extent is the induction of an ectopic neural plate by a graft comparable to the events of normal neural plate development? To address this, we used three complementary approaches. First, we explored the spatial and temporal patterns of expression of many components of the network during normal embryo development. Then, we conducted single cell RNAseq analysis of stages of neural plate development concentrating on these network components and their temporal hierarchical organization. Finally, we tested the activity of several of the identified putative regulatory elements in normal embryos to assess whether their activity in the neural plate *in vivo* resembles the hierarchy revealed by node grafts.

### Spatiotemporal expression of GRN components during normal neural plate development

A previous study identified genes whose expression differs between EGKXII-XIII (pre-primitive-streak) epiblast, neural plate at HH6-7 and non-neural ectoderm (Trevers et al., 2018). When these embryonic tissues (Figure 5A) are compared to neural induction, we find that the pre-streak epiblast is most similar to induced ectoderm after 5h of a node graft, while 9h and 12h time points are more closely related to mature neural plate (Figure 5B). This encouraged us to generate a detailed spatiotemporal map of transcripts during embryonic neural plate development. Genes that are differentially expressed in induced tissues (using a ±1.2 log_2_ fold change compared to uninduced in RNAseq, *p*-value < 0.05) were selected. *In situ* hybridization was performed for genes encoding 174 transcriptional regulators (including 123 that are represented in the GRN) at 4 stages: pre-streak (EGKXII-XIII), primitive streak (HH3-4), head process/early neural plate (HH5-6) and neural fold/tube (HH7-9).

**Figure 5.**
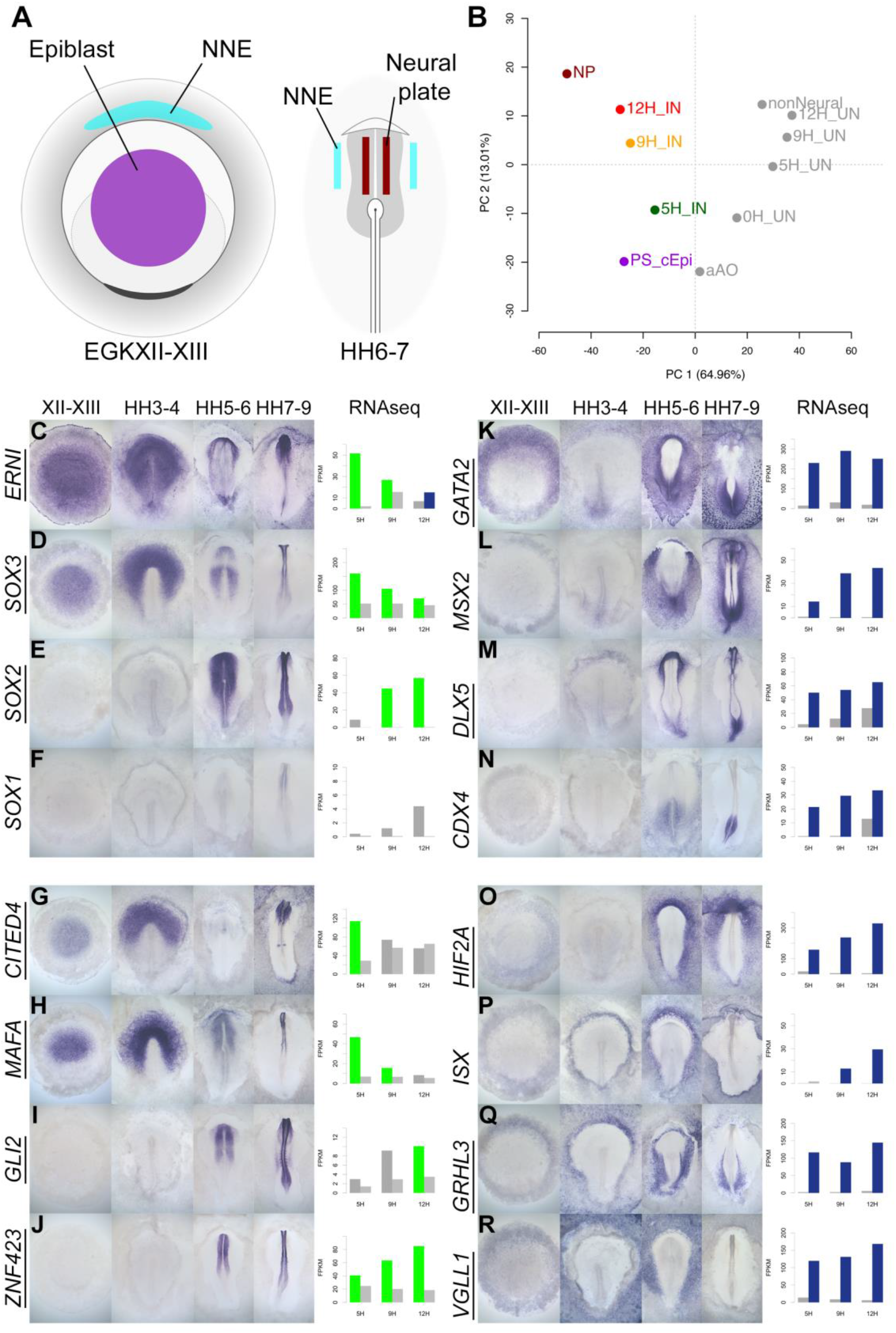
A spatiotemporal map of the normal embryo for transcripts differentially expressed during neural induction. (A) Central epiblast from pre-primitive-streak stage embryos (PS_cEpi) at EGKXII-XIII, neural plate (NP) at HH6-7 and corresponding non-neural ectoderm were dissected and processed by RNAseq. (B) Principal component analysis comparing prospective neural and non-neural tissues from the normal embryo to Induced (IN) and Uninduced (UN) ectoderm 5, 9 or 12h after a node graft. (C-R) *In situ* hybridization of genes encoding transcriptional regulators at 4 stages of embryonic development (EGKXII-XIII, HH3-4, HH5-6 and HH7-9) compared to their RNAseq expression after 5, 9 and 12h of a node graft. Bar charts plotted RNAseq values (FPKM) in Induced and uninduced tissues. Bars are shaded green when genes are upregulated in induced tissue (FPKM >10 in induced tissue and FC > 1.5 compared to uninduced). Bars are shaded blue when genes are downregulated (FPKM >10 in uninduced tissue and FC < 0.5 compared to uninduced). When a gene is neither upregulated nor downregulated, the bars are coloured in dark grey in induced and light grey in uninduced. Genes represented within the neural induction GRN are underlined.

There is a striking correspondence between neural induction by a node graft and neural development in terms of the spatiotemporal patterns observed. In addition to confirming the expression of known neural markers (*ERNI*, *SOX3*, *SOX2*, and *SOX1*; Figure 5C-F), the predictions from node grafts uncover many novel responses such as *CITED4* (*ERNI*-like), *MAFA* (*SOX3*-like), *GLI2* and *ZNF423* (*SOX2*-like) (Figure 5G-J). In total, 84/89 transcriptional regulators that are induced by a node graft were detected in neural tissues at some stage, 79 of which had not previously been associated with neural induction (Figure 5 and Suppl File S3). Only 5 genes (*HEY1*, *RFX3*, *STOX1*, *TAF1A* and *ZIC2*) could not be detected by *in situ* hybridization - these are induced at very low FPKM levels according to RNAseq. Likewise, a high proportion (65/85; 76%) of transcripts that are downregulated by a node graft are depleted in neural tissues relative to non-neural territories. Among these are known non-neural markers such as *GATA2* and *MSX2* and the extraembryonic marker *HIF2A* (Figure 5K, L, O). Neural/non-neural border markers (such as *DLX5*, *MSX1* and *TFAP2C*; Figure 5M and Suppl File S3) were also downregulated, supporting suggestions that neural induction involves the repression of an early border identity (Trevers et al., 2018). We also uncover several other genes (such as *ISX*, *GRHL3* and *VGLL1*) that are excluded from neural tissues (Figure 5P-R).

Importantly, this approach reveals quite subtle temporal changes in expression. For example, *CITED4* is sharply upregulated after 5h of a node graft but is later downregulated (Figure 5G). In the embryo, it is expressed in pre-streak epiblast and prospective neural plate at HH3-4, but these transcripts are then excluded from the neural plate at HH5-6. *CDX4* and *HOXA2* are initially repressed by a node before their expression levels increase at 12h – just as they become enriched in the posterior neural tube from HH7-9 (Figure 5N and Suppl File S3). These observations confirm that transcriptional responses to a node graft closely follow the events that occur during development of the embryonic neural plate.

### The GRN describes a temporal hierarchy occurring in single cells during neural development

To quantify the expression of multiple genes during normal neural plate development and to compare this to the temporal hierarchy of our GRN, we assessed the expression of network components in individual cells at different stages of normal neural development. Single cell RNAseq (scRNAseq) was performed on ectoderm dissected from chick embryos at HH4, HH6, HH8 and HH9+. These broad regions included prospective or mature neural plate and neural tube as well as non-neural ectoderm and neural plate border (Figure 6A).

**Figure 6.**
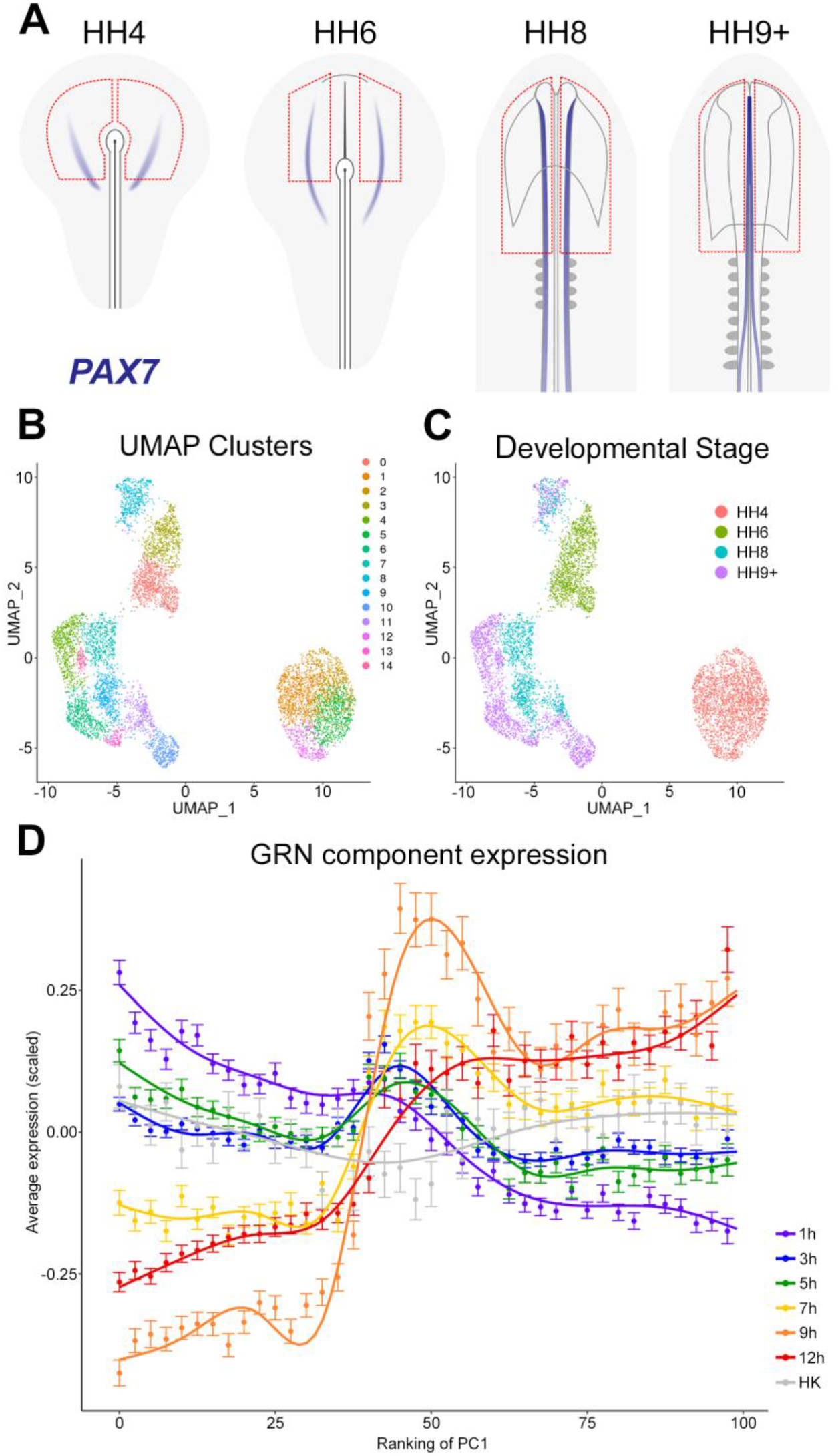
The GRN describes a temporal hierarchy occurring in single cells during neural development. (A) Broad regions of prospective neural plate, neural plate border and non-neural ectoderm were dissected at HH4, HH6, HH8 and HH9+ and processed for scRNAseq. *PAX7* expression marks the border between neural and non-neural tissues. (B) UMAP displaying unbiased clustering of cells into 15 clusters (0-14) representing distinct cell identities. (C) UMAP displaying the developmental stage from which the cells were collected; cells from HH4 and HH6 form independent clusters, whereas HH8 and HH9+ are more intermixed. (D) Temporal hierarchy of expression of GRN components in single cells during neural plate development. Prospective neural plate and neural tube cells (clusters 0, 1, 2, 5, 6, 7, 9, 13 and 14) were selected and ranked according to PC1 (see Figure S4C). The average gene expression (mean +/- SE) is plotted for groups of components that are induced at each time point in the GRN, compared to housekeeping genes (HK). A general additive model was fitted to visualize the smoothed expression profile across the ranking of PC1 (using bins of 2.5).

The non-linear dimensional reduction method UMAP was used to visualize unbiased cell clustering in low dimensionality space (Figure 6B). Cells collected from the two earliest developmental stages (HH4 and HH6) each form a distinct group, whereas cells from later stages (HH8 and HH9+) are clustered primarily according to cell type (Figure 6C). Cell identities were assigned to the 15 clusters using well-established markers (Figure S4A). This revealed clusters corresponding to placodal (cluster 8), neural crest (clusters 10, 11), and neural tube identities (clusters 4, 6, 7, 9, 13, 14) within the HH8 and HH9+ cell populations. At HH6, neural plate cells expressing *LIN28B*, *SETD2* and *SOX2* (cluster 0) are distinct from non-neural plate progenitors (cluster 3), which express *GATA2*, *DLX5* and *TFAP2A*. At HH4, prospective neural cells express *MAFA*, *ING5* and *LIN28B* (clusters 1, 2 and 5) whereas cluster 12 expresses the node marker ADMP.

To identify markers that are co-expressed in a cell-type specific manner, gene module analysis was performed using Antler (Delile et al., 2019). For a module to be retained for further consideration, we applied the criterion that at least 50% of its genes must be differentially expressed (logFC > 0.25, FDR < 0.001) in at least one cell cluster relative to the rest of the dataset. This condition defines seventeen distinct modules (Figure S4B), which support our cell type classifications. Module-3 shows co-expression of *PAX2*, *WNT4* and *NKX6-2* and is upregulated in prospective midbrain clusters 9 (at HH8) and 6 (at HH9+). Module-10 is defined by co-expression of *PAX7*, *SOX8*, *SOX10* and *TFAP2B* and is associated with neural crest (clusters 10, 11). Module-25 has low levels of expression of *GATA2* and *ID3* (clusters 1, 2, 5, 12) at HH4, which are then further downregulated at later stages, except in non-neural cells at HH6 (cluster 3) and placodal cells at HH8 (cluster 8).

To validate the temporal hierarchy of GRN components, their expression was assessed in prospective neural plate and neural tube cell populations (clusters 0, 1, 2, 4, 5, 6, 7, 9, 13, 14). Within this “neural” population, principal component (PC) analysis revealed that PC1 inversely correlates with developmental stage (Figure S4C), therefore cells were subsequently ranked according to their position along PC1 to explore expression dynamics. To visualize the expression of GRN components, genes were grouped according to the time point at which they are first expressed after a node graft; their average (normalized and scaled) expression was then plotted (Figure 6D). This reveals that components upregulated within 1, 3 or 5h of a node graft are initially highly expressed in prospective neural cells and that their expression decreases progressively across PC1. GRN components that are induced after 7, 9 or 12h are expressed at higher levels later, corresponding to maturing neural plate and neural tube. Taken together, our single cell RNAseq analysis uncovers the changes in gene expression accompanying development of the normal neural plate. Within this, GRN components defined from node graft experiments are appropriately expressed, suggesting that the temporal hierarchy of the neural induction assay closely replicates normal neural plate development.

### Responses to neural induction reveal enhancers that drive neural gene expression during normal development

Do the regulatory elements predicted by the GRN generated from Hensen’s node grafts correspond to those that regulate normal development of the neural plate in the embryo? To help define the putative enhancers more precisely, we combined the histone profiling described in the previous sections with ATACseq (Buenrostro, Giresi, Zaba, Chang, & Greenleaf, 2013; Buenrostro, Wu, Chang, & Greenleaf, 2015; Corces et al., 2017) to identify regions of open/active chromatin from induced and uninduced tissues after 5, 9 and 12h after a node graft. These tissues were then compared to ChIPseq (for H3K27ac/me3) and ATACseq performed on pre-primitive-streak epiblast and mature neural plate at HH6-7.

We chose 6 putative enhancers of GRN components for more detailed exploration. These were identified within the previously described CTCF loci using the overlapping combination of H3K27ac marks and ATACseq reads to predict enhancers that are likely to be active (Figures S5-S7). Each region was cloned into the pTK-EGFP vector (Uchikawa et al., 2003) to drive GFP expression via a minimal promoter; the reporter was then electroporated into the pre-primitive-streak stage (EGKX-XI) epiblast or the prospective neural plate at HH3/3+ (Figure 7A) and embryos cultured for 5h or to the desired stages.

**Figure 7.**
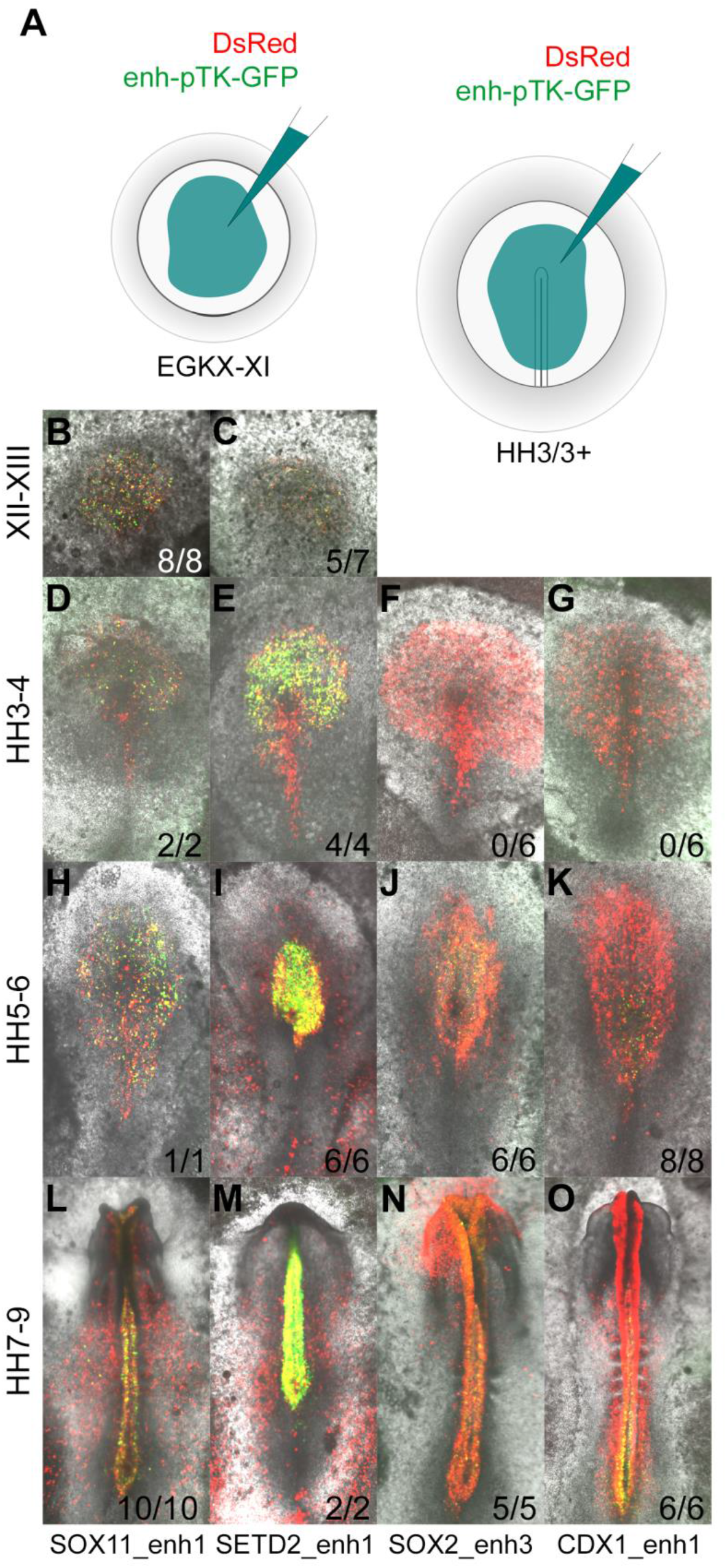
Enhancers identified by the neural induction assay have activity consistent with the expression of the gene during normal neural plate development. (A) Putative enhancers were cloned into the E-pTK-EGFP vector and co-electroporated into a broad region of the pre-streak stage (EGKX-XI) or HH3+ epiblast together with DsRed. (B-C) Activity of SOX11_enh1 and SETD2_enh1 at EGKXII-XIII; (D-G) Activity of SOX11_enh1, SETD2_enh1, SOX2_enh3 and CDX1_enh1 at HH4; (H-K) HH5-6; (L-O) HH7-9.

All 6 enhancers drive neural-specific GFP expression in a spatiotemporal pattern strikingly similar to the genes that lie in their proximity (Figures 7B-O and S5-S7). For example, SOX11_enh1 and SETD2_enh1 are enriched for H3K27ac in induced tissues after 5h (Figure S5A-B). Both are active in the epiblast at EGKXII-XIII and prospective neural plate at HH3-4, when *SOX11* and *SETD2* are expressed (Figure S5C-AY). SOX2_enh3 is “poised” after 5h of neural induction, but acquires an active signature at 9h when it gains H3K27ac and ATAC reads (Figure S6A). As expected, this enhancer is not active in the prospective neural plate at HH4 (Figure S6C-H); GFP positive cells are detected later when *SOX2* is expressed in the neural plate proper at HH5-6 (Figure S6I-N). Likewise, GLI2_enh1 and SIX3_enh2 also become active after 9h; GLI2_enh1 is detected throughout the neural plate, whereas SIX3_enh2 is restricted to the forming forebrain at HH6 (Figure S7A-Z). These five enhancers remain active in the neural tube in correspondence with H3K27ac marks and ATAC reads that persist after 12h of a node graft.

CDX1_enh1 is also active in the caudal neural tube at HH7-9, when it also weakly labels some presomitic mesoderm cells. Its activity is detected at HH5-6, but only in a few cells of the caudal lateral epiblast adjacent to Hensen’s node and not in the neural plate (Figure S6U-AL). This sparse activity overlaps closely with the discrete region from which neuro-mesodermal precursors, which are known to express *CDX1* (Gouti et al., 2017), are thought to contribute to both neural and somite tissues (Brown & Storey, 2000; Tzouanacou, Wegener, Wymeersch, Wilson, & Nicolas, 2009). Since the *CDX1* gene is expressed earlier (HH3-4), but only in mesoderm and primitive streak, the activity of this enhancer is likely to be associated with NMPs.

Taken together, these comparisons between node grafts and normal embryos strongly suggest that histone profiles that change in response to a node graft (upon which our network is based) can predict regulatory elements relevant to neural development. This establishes our GRN as a robust tool to describe the genetic hierarchy associated with the acquisition of neural identity.

### Browser resources

In addition to the BioTapestry GRN and other associated data described above, the results of this project, including data from node grafts in time course and from normal neural plate development, are made available on the UCSC Browser at https://genome.ucsc.edu/s/stern_lab/Neural_Induction_2021. This incorporates RNAseq, ChIPseq and ATACseq for all expressed and repressed genes (not limited to GRN components) and the annotated putative regulatory regions with predicted binding sites for GRN components. Finally, to help to predict potential conservation of the regulatory sites across species, we performed DREiVe analysis (Khan et al., 2013; Sosinsky, Honig, Mann, & Califano, 2007; Streit et al., 2013) for domains within a 500kb window around each of the genes represented in the GRN, using human (hg38), mouse (mm10), rat (rn6), golden eagle (aquChr2), zebrafish (danRer10) and chicken (galGal5). Blocks of sequence with conserved motifs are included as a track in the browser.

## Discussion

This study unfolds the full complexity of the responses to signals from the “organizer” in very fine time-course up to the time of appearance of mature neural plate markers, such as *SOX1*. We show that the hierarchy of responses to a graft of Hensen’s node into a remote part of the embryo closely recapitulates the events of normal neural plate development, including appropriate activity of enhancers identified based on the graft assay. Our GRN, comprising 175 transcriptional regulators and 5,614 putative regulatory interactions between them, allows prediction of the consequences of manipulating the activity of these components. This represents the most comprehensive example to date of a network modelled on both transcriptional and chromatin changes that occur in the embryo. These changes were selected in an unbiased way by comparing induced and uninduced tissues to identify events specific to neural induction. By following this process over very short intervals (1-3h), we can predict interactions likely to occur via direct transcription factor binding events. This is accompanied by a rich resource (deposited in the UCSC genome browser) that includes comprehensive genome-wide datasets in time-course for all transcribed loci, epigenetic marks and annotations for prediction of evolutionary conservation of the identified enhancer elements. This is the first comprehensive description of the progression of molecular events of neural induction in any species, and the first GRN for this process.

Is it possible to identify specific changes associated with the key concepts of developmental biology, such as “induction” and “commitment” (Slack, 1991; Stern, 2004) to a neural fate? These concepts have classically been defined based on specific embryological manipulations. Based on timed organizer grafts followed by their removal, a critical time was established at 12-13 hours after a graft (Gallera, 1965, 1971a, 1971b; Gallera & Ivanov, 1964), using similar conditions to those of our node grafts here. This represents the endpoint of our GRN which coincides approximately with the onset of expression of the definitive neural tube marker *SOX1* (Pevny, Sockanathan, Placzek, & Lovell-Badge, 1998).

Previous functional studies have established that competent ectoderm cannot respond to BMP-inhibitors like Chordin unless it has previously been exposed to a graft of the organizer for at least 5 hours (Streit et al., 1998). This led to the identification of a few genes whose expression is associated with this transition (Gibson et al., 2011; Costis Papanayotou et al., 2013; Pinho et al., 2011; Sheng et al., 2003; Streit et al., 2000), but the present experiments greatly expand and enrich this by placing genes in the context of a comprehensive network of gene interactions. Future experiments can take advantage of this information to determine the mechanisms that cause epiblast cells to become sensitive to BMP inhibition at this time point, and the upstream signals responsible. For the time being we feel that functional definitions of the developmental biology concepts will remain key to defining the important molecular changes that underlie these properties of developing tissues, especially given the complexity of the changes in gene expression accompanying neural induction.

Our analysis reveals thousands of sites across the genome whose marks change with the progression of neural induction. Each locus contains genes whose expression changes during this cascade and several putative regulatory elements. As previously described for other systems such as embryonic stem cells in culture, regulatory elements may be decorated simultaneously by marks associated with activation (e.g. H3K27ac) and with repression (e.g. H3K27me3); these have been called “poised” enhancers (Creyghton et al., 2010; Cui et al., 2009; Heintzman et al., 2009; Xu et al., 2009) and often interpreted to mean that the site is in transition from one state to the other, or perhaps able to go either way. Our fine time course certainly reinforces this concept; in many cases we observe that cells pass from one state (e.g. H3K27me3 without acetylation) to the opposite, passing through a “poised” state at the intermediate time point. This could suggest that these changes can occur sequentially in a cell. However, it is also possible that this progression can reflect cells within a tissue that do not undergo the change synchronously, but some cells start earlier than their neighbours, the tissue gradually becoming more homogeneous. This sort of progression is sometimes seen by in situ hybridization or time-lapse movies of reporter activity (for example to visualize the activity of enhancer N2 associated with the SOX2 gene (C. Papanayotou et al., 2008; Uchikawa et al., 2003). Most likely these “poised” sites reflect a combination of these two reasons.

Among the many regulatory elements predicted by our analysis are some that have been described previously (Iida et al., 2020; Okamoto et al., 2015; Uchikawa et al., 2003; Uchikawa et al., 2011). Their analysis was based on nested deletions followed by testing the activity of the fragments in a vector containing a heterologous minimal promoter. In our study we initially predicted the elements based on a combination of marks and regions marked by ATACseq as being “open”. In some cases, there are differences in the length of the sequence considered to act as an enhancer by both studies. For example, our SOX2_enh3 partially overlaps with, and its behaviour upon electroporation mimics the activity of, the previously identified *SOX2* D1 enhancer (Iida et al., 2020; Okamoto et al., 2015). Therefore, our ChIPseq analysis can verify known enhancers as well as identifying many new candidates.

In conclusion, our study provides the first GRN to represent the changes that accompany the process of neural induction after grafting an organizer to an ectopic site, and how it relates to normal neural plate development, in fine time course. We also provide a comprehensive resource including not only the network components (transcriptional regulators with predicted cross-regulatory interactions), but also a genome-wide view of all transcriptional changes, H3-K27 acetylation/methylation and chromatin conformation (ATACseq) during this process, from the time cells are initially exposed to neural inducing signals to the time at which embryological experiments suggest that the tissue has become “committed” to a neural identity. We envisage that the GRN and the accompanying resource will be invaluable tools to study other important aspects of early neural development, such as the signalling inputs responsible for these changes, the molecular basis of the competence of cells to respond to neural inducing signals at different locations and stages, and aspects of neural patterning leading to regional diversification of the central nervous system. To aid in cross-species comparisons and possible studies on evolutionary changes that have occurred in neural induction, the browser resource also includes the results of computational prediction of conservation of the key GRN loci among several vertebrate species, using DREiVe, a computational tool designed to identify conserved sequence motifs and patterns rather than precise sequence conservation (Khan et al., 2013; Sosinsky et al., 2007; Streit et al., 2013).

## MATERIALS AND METHODS

### Chicken eggs and embryo culture

Fertile hens’ eggs (Brown Bovan Gold; Henry Stewart & Co., UK), were incubated at 38°C in a humidified chamber to the desired stages. Embryos were staged according to (Hamburger & Hamilton, 1951) in Arabic numerals or (Eyal-Giladi & Kochav, 1976) in Roman numerals for pre-primitive streak (pre-streak) stages. Embryos were harvested and dissected in Pannett-Compton saline (Pannett & Compton, 1924) or 1x PBS, before being fixed in 4% paraformaldehyde (PFA) overnight at 4°C or prepared for *ex ovo* culture. All experiments were conducted on chicken embryos younger than 12 days of development and therefore were not regulated by the Animals (Scientific Procedures) Act 1986.

Chicken embryos were cultured *ex ovo* using a modified New culture method (New, 1955; Stern & Ireland, 1981) as previously described (Voiculescu, Papanayotou, & Stern, 2008). In brief, eggs were opened and the thin albumin was collected. Intact yolks were transferred to a dish containing Pannett-Compton saline. The vitelline membrane was cut at the equator while keeping the embryo central. The membrane was then slowly peeled away from the underlying yolk to keep the embryo attached. Membranes were placed on a watch glass with the embryo orientated ventral side up. A glass ring was positioned over the embryo and the edges of the membrane wrapped over the ring. This assembly was then lifted out of the dish and adjusted under a dissecting microscope while keeping the embryo submerged in a small pool of saline. The vitelline membrane was gently pulled taut around the glass ring before the excess was trimmed. Excess yolk was cleared from the membrane and embryo by gently pipetting a stream of saline and then replaced with fresh liquid, leaving the embryo on an optically clear membrane. At this point, embryos can be cultured as they are or with the addition of a node graft which were transferred to the ring and positioned while the embryo is submerged. To complete the culture, saline was removed from around and within the ring without disturbing the embryo and grafts. The dry ring was then transferred to a 35mm Petri dish containing a shallow pool of thin albumin. The edges of the ring were pressed down to prevent it from floating, leaving the embryo supported by a shallow bubble of albumin beneath the membrane. Completed cultures were incubated in a humidified chamber at 38°C for the desired length of time. After culture, embryos were fixed on the membrane with 4% PFA or submerged with ice-cold saline to allow tissue dissection.

### Neural induction assays

Neural induction assays were performed as previously described (Stern, 2008; Streit & Stern, 2008). For Hensen’s node grafts, chick donors and hosts at Hamburger Hamilton (HH) 3+/4- were used and New cultures incubated for 1, 3, 5, 7, 9 or 12h at 38°C in a humidified chamber. One or two nodes were grafted per embryo, placed contralaterally within the inner third of the area opaca, at or above the level of the host node. This region is competent to respond to neural inducing signals but only contributes to the extra-embryonic membranes and not the embryo proper (Streit et al., 1997). Nodes were grafted with their endodermal surface in contact with the host epiblast.

### RNAseq tissue collection and processing

For RNAseq, HH4-chick nodes were grafted to the area opaca of HH4-chick hosts. A single node was grafted to the left or right side per host, and embryos were cultured for 5, 9 or 12h. Node grafts were removed by submerging the embryo in saline while still attached to the membrane. Fine syringe needles (27G or 30G, B D Microlance) were used gently lift the grafted tissue away from the underlying epiblast. Where grafts were firmly attached, 0.12% trypsin dissolved in saline was gently pipetted over the graft site to assist in removal. After dissection, trypsin activity was neutralized by treating embryos briefly with heat-inactivated goat serum (Stern, 1993) which was removed by washing with fresh saline. The induced epiblast directly beneath the graft, which appeared greyish and thickened, was dissected using syringe needles and mounted insect pins. Uninduced epiblast tissue from same position on the contralateral side was also dissected. Tissue samples per condition were pooled, frozen on dry ice and stored at -80°C. At each time point (5, 9 or 12h) a total of 50 induced and contralateral uninduced pieces of tissue were collected. A further 50 pieces of uninduced tissue were collected from HH4-embryos that had not been cultured, representing a 0h control. Tissue samples for each condition were then pooled and lysed in 1mL TRIzol (Invitrogen) for RNA extraction.

Transcriptome sequencing was conducted by Edinburgh Genomics (formerly ARK Genomics; Roslin Institute, University of Edinburgh). Total RNA was extracted and the quality was assessed using the Agilent 2100 Bioanalyzer -all samples registered a RIN value between 9.0-10.0. Libraries were constructed using the Illumina® TruSeq mRNA library preparation kit. In brief, mRNAs were purified using oligo-dT conjugated magnetic beads before being chemically fragmented to on average 180-200bp. Fragments were then transcribed using short random primers and reverse transcriptase to produce single-stranded cDNA, from which double-stranded cDNA was generated by DNA polymerase I and RNase-H. After synthesis, cDNA was blunt-ended and a single A-base added to the 3’ end. Sequencing adapters were ligated via a T-base overhang at their 3’ and these libraries were then purified to remove unincorporated adapters before being enriched by 10 cycles of PCR. Library quality was checked by electrophoresis and quantified by qPCR. The 7 RNA libraries were sequenced over 2 lanes via 100-cycle, paired-end sequencing using the Illumina® HiSeq 2000 system.

### Whole-mount in situ hybridization

Antisense riboprobes were generated from cDNA plasmid templates or Chick Expressed Sequence Tag (ChEST) clones (Source Bioscience). Templates were linearized by restriction digest (Suppl Table S10) or by PCR (Suppl Table S11) using M13 forward (5’-GTAAAACGACGGCCAGT-3’) and M13 reverse (5’-GCGGATAACAATTTCACACAGG-3’) primers (Suppl Table S12). The products were transcribed using SP6, T7 or T3 (Promega) and DIG-labelled nucleotides. DNA templates were digested using RNase-free DNase and the RNA probe was precipitated and resuspended in water. Working probes were diluted to 1μg/mL in Hybridization (HYB) buffer and stored at -20°C.

Whole-mount in situ hybridization was performed as described previously (Stern, 1998; Streit & Stern, 2001), but omitting pre-adsorption of the anti-DIG-AP antibody. Stained embryos were imaged from the dorsal perspective using an Olympus SZH10 Stereomicroscope with an Olympus DF PlanApo 1X objective and an Olympus NFK 3.3x LD 125 photo eyepiece. Images were captured using a QImaging Retiga 2000R Fast 1394 camera and QCapture Pro software as 24-bit, color TIFF files at 300dpi with dimensions of 1600 x 1200 pixels.

A spatiotemporal expression map was generated for genes that are differentially expressed after either 5, 9 or 12h of a node graft. These were selected from the *galGal3* and *galGal4* RNAseq analyses according to the following criteria. In *galGal3*, genes that are upregulated by a log_2_ fold change of ≥1.2 or downregulated by ≤-1.2 in either Cufflinks or DESeq analyses, and were statistically significant using a p-value of ≤0.05 (i.e. coloured red or orange in Suppl Table S15-16) were selected. In *galGal4*, genes that were upregulated by a log_2_ fold change of ≥1.2 or downregulated by ≤-1.2 in DESeq and were statistically significant using a p-value of ≤0.05 (i.e. coloured pale blue in Suppl Table S17-18) were selected. In situ hybridization was performed for transcriptional regulators that satisfied these criteria (Suppl Table S19).

### NanoString tissue collection and processing

Uninduced tissue or induced epiblast that had been exposed to node grafts was dissected as previously described. Four to eight pieces of tissue were collected in triplicate per condition. Tissues were dissected in ice-cold 1xPBS and collected on ice. All excess solution was removed using a fine needle and promptly processed by adding a total of 1µL of lysis buffer from the RNAqueous-Micro Total RNA Isolation kit (ThermoFisher). Tubes were immediately snap-frozen on dry ice and stored at -80°C.

NanoString experiments were run on the nCounter Analysis System using a custom codeset and following NanoString guidelines. The codeset consisted of 386 probes (Suppl Table S8) belonging to various categories. Transcriptional regulators that are upregulated (coloured red) at 5, 9 or 12h with log_2_ scaled fold change (FC) ≥1.2 and an induced base mean ≥45 were selected from DESeq analysis using *galGal3* and *galGal4* assemblies. Transcriptional regulators that are downregulated (coloured green) at 5, 9 or 12h with log_2_ scaled FC ≤-1.2 and uninduced base mean ≥200 were also selected. Other genes that respond to 5h exposure to a node graft (Gibson et al., 2011; Costis Papanayotou et al., 2013; Pinho et al., 2011; Sheng et al., 2003; Streit et al., 2000) were also included. Probes were also designed against housekeeping genes (ACTB, GAPDH), markers of apoptosis and proliferation, transcriptional readouts of FGF, BMP, WNT, Retinoic acid, Notch and Hedgehog signalling; epithelial, mesodermal, endodermal, neural plate border, pre-placodal, neural crest and Hensen’s node markers, and standard positive and negative NanoString controls.

Tissue samples were processed using the NanoString master kit and following manufacturer’s instructions. Probes were hybridized to lysates overnight for 17h at 65°C before the reactions were transferred to the NanoString prep-station robot and processed using the “high-sensitivity” program. Probe-target complexes were digitally counted from 600 fields of view on the NanoString Analyzer.

### Enhancer cloning and electroporation

Enhancers were identified based on changing active, inactive or poised chromatin signatures over 5, 9 and 12h of neural induction. These were referenced against the 5, 9 and 12h ATACseq data to select putative enhancers with accessible chromatin. The primers used for cloning are detailed in Suppl Table S13.

Enhancers were cloned from chick gDNA as previously described (Chen & Streit, 2015). The correct sized fragments were amplified by PCR using PCRBio Ultramix (PCRBiosystems) and ligated into the pTK_JC_EGFP plasmid (Chen & Streit, 2015) using T4 ligase (NEB). Ligation products were transformed into DH5α *E.coli* and grown on agar plates with Ampicillin selection. Clones were screened by colony PCR using PCRBio Ultramix and pTK_Fwd (5’-GTGCCAGAACATTTCTCTATCG-3’) and pTK_Rev (5’-GTCCAGCTCGACCAGGATG-3’) primers. Positive clones were cultured overnight in 5mL LB broth plus Ampicillin and the constructs were purified by mini-prep (Qiagen). Enhancer inserts were sequenced using Citrine_Fwd (5’-TGTCCCCAGTGCAAGTGC-3’) and Citrine_Rev (5’-TAGAACTAAAGACATGCAAATATATTT-3’) primers. Plasmids with sequence verified inserts were maxicultured in 200mL, of which 1mL used to make a 50% glycerol stock and stored at -80°C. Bulk plasmid DNA was extracted using the Endofree Plasmid Maxi-prep kit (Qiagen) following manufacturer’s instructions and resuspended in 100µL of sterile water. Further purification was conducted to reduce residual salts which interfere with electroporation efficiency and embryo damage. Samples were spun for 15mins/15,000rpm/4°C to pellet any particulates. The supernatant was removed to a fresh tube, to which 10µL of 3M NaOAc pH5.2 and 250µL of 100% EtOH was added to precipitate the plasmid DNA. This was pelleted by centrifugation at 15mins/15,000rpm/4°C and the pellet was washed with 70% EtOH and spun again for 10mins/15,000rpm/4°C. The supernatant was removed and the pellet allowed to air-dry completely, before being resuspended in 50µL of sterile water and the concentration checked by nanodrop and adjusted to 5mg/mL.

Enhancer plasmids were co-electroporated with pCMV-DsRed-Express-N1 (Clontech) which labels all electroporated cells. Tissues were electroporated using an anode electroporation chamber and wire cathode according to (Voiculescu et al., 2008) with minor modifications. Electroporation mix was made fresh by combining the following in water: enhancer_TK_EGFP plasmid (2mg/mL), pCMV-DsRed-Express-N1 (1mg/mL), 6% sucrose, 0.02% Brilliant Blue FCF. This was backloaded into a capillary needle with a fine tapered tip. The anode chamber was filled with 1x PBS and the electrodes positioned ≈2cm apart.

Harvested embryos were positioned directly between the electrodes with the dorsal side (epiblast) upwards, closest to the cathode. Electroporation mix was applied using a mouth aspirator attached to the capillary needle, and by bringing the needle tip in close contact with the embryo. For pre-streak EGKX-XI embryos, electroporation mix was applied broadly across the central epiblast region and immediately electroporated using the following current settings: 3.5-4.0V, 5 pulses, 50ms duration and a 100ms gap. At HH4-stages, electroporation mix was applied broadly across the prospective neural plate and primitive streak and immediately electroporated using 4.5-5V, 5 pulses, 50ms duration and 100ms gap. After electroporation embryos were transferred to vitelline membranes, prepared for New culture and cultured to the desired stages (Voiculescu et al., 2008).

Enhancer activity was imaged using a Leica SPEinv microscope with a 5x objective and LAS-X software. Images were captured at 2048x2048 pixels, speed 400ms, and 1x zoom with an averaging of 4. EGFP was excited using a 488nm laser and detected with a 492-543nm wavelength filter. DsRed was excited using a 532nm laser and detected with a 547-653nm wavelength filter. Files were saved as .lif format and composite images were stitched together in Fiji (Schindelin et al., 2012) using the Pairwise Stitching plugin (Preibisch, Saalfeld, & Tomancak, 2009).

### ChIPseq tissue collection and processing

In brief, tissues were dissected in ice cold 1x PBS and transferred to low binding PCR tubes containing 50-100µL of 1x Protease Inhibitor Cocktail in 1xPBS (cOmplete mini EDTA-free; Roche) on ice. Where necessary, 0.12% trypsin or 1XPBS + 200mM EDTA was used to aid dissection. Treated tissues were rinsed with PBS to remove residual trypsin. Tissues were collected in batches every 30mins before tubes were spun at 100G/3min/4°C to gently pellet the tissue. Tubes were snap frozen in liquid nitrogen before storage at -80°C for up to 6 months. For pre-streak EGK XII-XIII embryos, the hypoblast was removed and central epiblast was collected from 6 embryos in triplicate. After removal of the underlying mesoderm and endoderm, the neural plate at HH6-7 was harvested from 20 embryos in triplicate. For 0, 5, 9 and 12h node induced and contralateral uninduced area opaca, 14-30 pieces of tissue were collected in triplicate. A further 60 pieces of 9h and 12h uninduced tissue were collected and processed separately after it was discovered that 30 pieces of uninduced tissue at these time points did not generate enough material for ChIP.

ChIP was performed by micrococcal nuclease (MNase) digestion to fragment the chromatin following a previous protocol (Brind’Amour et al., 2015), followed by immunoprecipitation according to the Low Cell ChIPseq kit (Active Motif, 53084) with some modifications. Corresponding induced and uninduced samples were processed side-by-side, each as 3 reactions: H3K27ac, H3K27me3 and input. In brief, 200µL of protein G agarose beads were added to 2x 1.5mL low binding tubes and spun for 3mins at 1250*g* and 4°C. The supernatant was removed and the beads were washed with 690µL of TE buffer at pH 8.0. These tubes were spun again for 3mins at 1250*g* and 4°C and the supernatant removed before bead blocking reactions were set up. To one tube, 200µL of TE buffer pH 8.0, 20µL of blocking reagent AM1 and 20µL of BSA were added. This tube of “pre-clearing” beads was incubated for 3-6h on a 4°C rotator. To the second tube, 180µL of TE buffer pH 8.0, 20µL of blocking reagent AM1, 20µL of BSA and 20µL of Blocker were added. This tube of “IP” reaction beads was incubated overnight on a 4°C rotator.

While beads were blocking, tissues were prepared for nuclei isolation and MNase digestion. A pair of samples (induced and contralateral uninduced) were thawed on ice for 5-10mins and the tissues pooled into one tube per condition using low binding tips. Pooled samples were spun at 100G/3mins/4°C and the excess supernatant was removed without disturbing the tissues, to leave ≈5µL remaining. To each tube, 20µL of Nuclei lysis buffer (NLB) was added, containing 16µL of Nuclei isolation buffer (Sigma N3408), 2µL 1x protease inhibitor (Roche cOmplete mini, EDTA-free) and 2µL of 1% IGEPAL CA-630. Tubes were gently tapped to mix and incubated on ice for 5mins. Nuclei were dissociated gently by pipetting 15 or 30 times without making bubbles, then incubating samples on ice for 15mins before repeating these steps 4-5 times as listed in Suppl Table S14.

Chromatin was then fragmented using MNase. This was carried out in two steps to fragment the easily digestible chromatin and then re-digest the more compact chromatin in a second longer digestion. A 3x master mix containing 169.5µL of sterile H_2_O, 30µL of 10x MNase buffer (NEB), 30µL of 50% PEG 6000 and 6µL of 100mM DTT (made fresh) was prepared. In a separate tube, 3µL of 10x MNase buffer (NEB M0247) was diluted in 25µL of H_2_O, before adding 2µL of MNase, to give a 1:15 dilution of MNase enzyme. Of this, 4.5µL of MNase 1:15 was added to the 3x master mix, to give a total volume of 240µL. Then, 50µL of 3x MNase master mix was added to each tube containing 25µL of isolated nuclei. The mixture was pipetted 15 times to mix without making bubbles, while tubes were kept on ice. This mixture was transferred to a fresh 0.2mL low binding PCR tube at room temperature and the digestion incubated at room temperature for 2.5mins. This tube was returned to ice and 6µL of 200mM EDTA added. Tubes were spun at 500*g* and 4°C for 5min to pellet larger undigested chromatin and the supernatant (containing smaller, readily digestible chromatin) was transferred to a fresh 1.5mL low binding microfuge tube. A further 30µL of the MNase master mix was added to the remaining chromatin pellets on ice and resuspended by pipetting 60 times, before tubes were incubated for 6 minutes at room temperature. To break up remaining nuclear debris, 4µL of 200mM EDTA and 4µL of 1% TX100 + 1% Sodium deoxycholate were added and the mixture pipetted 30 times. All 38µL of this second digestion was collated with the first supernatant and kept on ice (total volume ≈110µL).

Once the “pre-clearing” beads were ready (after 3-6h) they were spun for 3mins/1250*g*/4°C. The supernatant was removed and the beads washed with 665µL of ChIP Buffer (from Active Motif kit) by inverting the tube several times. Beads were spun again for 3mins/1250*g*/4°C and supernatant removed again, before 232µL of ChIP buffer was added. An even suspension of 100µL of beads was added to each tube of MNase digested chromatin (≈110µL). Then, 10µL of proteinase inhibitor cocktail, 10µL PMSF (from the Active Motif kit) and a further 290µL of ChIP buffer was added, giving a total volume of 520µL. Tubes were then placed on a rotator at 4°C for 3h to pull down chromatin fragments that bind non-specifically to the protein G agarose beads.

After “pre-clearing”, samples were spun at 3500 RPM and 4°C for 3mins and the supernatant was divided into 3 separate 0.5mL low binding PCR tubes: 200µL each into H3K27ac and H3K27me3 tubes and the rest into the input tube (≈60µL). To the experimental tubes, 2µL of H3K27ac (Active Motif 39133, lot# 31814008) or H3K27me3 (Active Motif 39155, lot# 31814017) antibody was added, and these were incubated overnight at 4°C. To the input, 80µL of low EDTA TE buffer was added and stored overnight at 4°C.

The following day, ChIP reaction tubes were spun briefly at 4°C to collect the solution. “IP” reaction beads were removed from the rotator and mixed to an even suspension by pipetting with a wide bore tip. Keeping the beads well mixed, 50µL of bead slurry was added to each immunoprecipitation and the tubes returned to the 4°C rotator for 3-4h to pull down ChIPed chromatin. Next, samples were washed according to Section G of the Active Motif Low Cell ChIPseq manual and eluted with 100µL of Elution Buffer AM4.

Chromatin was de-crosslinked by incubating ChIP and input samples at 65°C for 2h. Then 125µL of Phenol:Chloroform:Isoamyl Alcohol 25:24:1 and 64µL of Chloroform:Isoamyl alcohol mixture was added and samples shaken vigorously for 15secs before centrifuging at 16,000*g*/15min/4°C. The top aqueous layer was transferred to a fresh low binding tube. An additional 100µL of nuclease free water was added to the original tube, which was shaken and centrifuged again. Then the aqueous top layer was removed and added to the first. To this, 2µL of blue glycogen and 20µL of 3M NaOAc pH5.2 were added and samples flicked to mix. Finally, 660µL 100% ethanol was added, and tubes flicked to mix well before precipitating overnight at -20°C. The next day, the chromatin was centrifuged at 16,000*g*/30min/4°C. Supernatant was removed and the pellet washed with 900µL of cold 70% EtOH. The pellet was spun at 16,000*g*/30min/4°C and washed with 70% EtOH again, a total of 4 times. Then, chromatin pellets were air-dried completely for 10-15mins and resuspended in 20µL of TE low EDTA buffer overnight at 4°C, before being stored at -20°C until library preparation.

### ChIPseq Library Preparation

Libraries were prepared using the Next Gen DNA Library kit (Active Motif 53216) and Next Gen Indexing kit (Active Motif 53264) but using ProNex® size-selective beads (Promega) to enrich for mono and di-nucleosomes (≈200-500bp). Each sample of ChIPed chromatin was thawed on ice, to which 20µL of low EDTA TE Buffer was added. For each library, “Repair 1” reaction mix was prepared according to the Active Motif manual by combining 13µL of low EDTA TE, 6µL of Buffer W1 and 1µL of Enzyme W2. This was added to the 40µL of chromatin and mixed by pipetting. Reactions were incubated for 10mins at 37°C in a thermocycler with the lid open. To clean up the reaction, 180µL (3x volume) of evenly suspended ProNex beads were added and incubated for 10mins at room temperature before placing on a magnetic rack for 5mins. The supernatant was removed and beads were washed 3 times with 100µL of wash buffer, for 30-60s each. The final wash buffer was removed and the beads were air-dried for 5mins. Next, Repair II reactions were prepared by combining 30µL of low EDTA TE, 5µL of Buffer G1, 13µL of Reagent G2, 1µL of Enzyme G3 and 1µL of Enzyme G4. This was added to dry beads and pipetted to mix. Tubes were incubated on a thermocycler for 20mins at 20°C with lid open. Next, 90µL (1.8x volume) of PEG NaCl was added to each reaction and mixed by pipetting 15 times. Samples were incubated at room temperature for 10mins and then on the magnetic rack for 5mins and beads washed and air dried as before. Next, Ligation I mix was prepared by combining 20µL of low EDTA TE, 3µL of Buffer Y1, and 2µL of Enzyme Y3, which was added to each tube of dry beads and mixed by pipetting. To each sample, 5µL of Y2 index was added to uniquely barcoded libraries. Reactions were mixed by pipetting 60x and incubated in a thermocycler for 15mins at 25°C with the lid open. To clean up, 39µL (1.3x) of PEG NaCl was added to each tube and mixed by pipetting 15x. Samples were incubated for 10mins at room temperature and 5mins on a magnetic rack, before beads were washed and air-dried as before. Ligation II mix was prepared by combining 30µL of low EDTA TE, 5µL of Buffer B1 2µL of Reagent B2-MID, 9µL of Reagent B3, 1µL of Enzyme B4, 2µL of Enzyme B5, and 1µL of Enzyme B6. This was added to the dried beads and mixed by pipetting 30x, before incubating in a thermocycler for 10mins at 40°C with the lid open. Then, 55µL (1.1x) of PEG NaCl was added and mixed by pipetting 30x. Samples were incubated for 10mins at room temperature and 5mins on a magnetic rack before beads were washed and air-dried as before. DNA was eluted by adding 50µL of elution buffer and pipetting 15x. Tubes were incubated tubes at room temperature for 5mins, and then on a magnetic rack for 5mins. The eluate was transferred to a clean tube and the beads washed with a further 10µL of elution buffer. This was collected and combined with the first 50µL of eluate.

DNAs were size-selected using 1.2x/0.4x ratios of ProNex beads to purify ≈200-500bp fragments. First, 72µL of ProNex beads were added to the eluate and mixed by pipetting 30x. Samples were incubated for 10mins at room temperature and for 5mins on a magnetic rack. The supernatant (containing fragments <500bp) was transferred to a clean tube. Then, a further 24µL of ProNex beads were added to this supernatant and mixed 30x by pipetting. This was incubated for 10mins at room temperature followed by 5mins on a magnetic rack. The supernatant was removed and beads (with DNA fragments >200bp bound) were washed 3 times with 100µL of wash buffer, each time soaking beads for 30-60s. The final wash buffer was removed and beads air-dried for 5mins. DNA was eluted by adding 10µL of low EDTA TE, pipetting 10x to mix and incubating for 10mins at room temperature. After 5mins on a magnetic rack, the eluate was transferred to a fresh tube and the beads incubated for a further 10mins with 10µL of TE. This was collected and combined with the first collection to give a total of 20µL.

Libraries were amplified by adding amplification mix containing 10µL of low EDTA TE, 5µL of Reagent 1, 4µL of Reagent 2, 10µL of Buffer R3 and 1µL of Enzyme R4 to each tube. This was mixed by pipetting and incubated on a thermocycler using the following conditions: Denature1-98°C for 30s, Denature2-98°C for 10s, Anneal- 60°C for 30s, Extension- 68°C for 60s, Repeat to Denature2 for total 13 cycles, Final extension- 68°C for 60s, Soak- 4°C forever.

Libraries were purified by adding 75µL (1.5x volume) of ProNex beads and pipetting 60x. Samples were incubated at room temperature for 10mins, and 5mins on a magnetic rack. The supernatant was discarded and beads washed 3 times with 100µL of wash buffer, before being air-dried for 5mins. Libraries were finally eluted in 20µL of TE and their quality checked by Tapestation. They were sequenced by 75bp single end reads to a depth of ≈20million reads per library using the NextSeq 500 system.

### ATACseq tissue collection and processing

OMNI-ATACseq was conducted according to (Corces et al., 2017) with some adjustments, which was in turn based on (Buenrostro et al., 2013; Buenrostro et al., 2015). In brief, 0, 5, 9 and 12h node induced and contralateral uninduced tissues were dissected in ice cold 1x PBS and transferred to low binding PCR tubes containing 1x Protease Inhibitor Cocktail in 1xPBS (cOmplete mini EDTA-free; Roche) on ice. Where necessary, 0.12% trypsin was used to aid dissection, and treated tissues rinsed with 1xPBS to remove residual trypsin activity. Tissues were collected in batches every 30mins before tubes were spun at 100*g*/3min/4°C to gently pellet tissues. Tubes were snap frozen in liquid nitrogen before storage at - 80°C for up to 4 months. At each time point, 14-22 pieces of tissue were collected in duplicate.

Once sufficient tissues were collected, they were thawed on ice and the nuclei isolated in 2mL of 1x Homogenization Buffer Unstable (HBU) with a Dounce homogenizer on ice. HBU 1x contains 5mM CaCl_2_, 3mM Mg(Ac)_2_, 10mM Tris pH7.8, 320mM Sucrose, 0.1mM EDTA, 0.1% NP-40, 0.02mM PMSF and 0.3mM β-mercaptoethanol, dissolved in H_2_O. Tissues were ground using 15 strokes with the loose pestle and 20 strokes with the tight pestle. Homogenate was then passed through a 30µm nylon mesh filter into a fresh 2mL low binding microfuge tube. Next, nuclei were gently pelleted by centrifugation for 10mins/500*g*/4°C and the supernatant removed. The pellet was resuspended in 200µL of 1xHBU by gentle pipetting. Nuclei were counted by taking 6x10µL homogenate samples, staining them with DAPI and viewing on a haemocytometer. After counting, nuclei were washed with 1mL of ATAC-RSB+0.1% Tween-20 and spun for 10mins/500*g*/4°C to pellet. The supernatant was carefully removed and the nuclei pellet was resuspended in 50µL of transposition mix by pipetting. The volume of Tn5 transposase (Illumina, 20034197) added to each reaction was scaled to 0.1µL per 1000 nuclei, so that ATAC samples with different numbers of nuclei were treated equivalently. Therefore, transposition mix contained; 25µL of 2x TD Buffer, 0.5µL of 1% Digitonin, 0.5µL of 10% Tween-20, 16.5µL of 1xPBS, XµL of Tn5 transposase (0.1µL of Tn5 per 1000 nuclei), topped up with sterile H_2_O up to 50µL. Transposition reactions were incubated for 30mins at 37°C in a thermomixer with 1000rpm shaking. Immediately after samples were purified using the Zymo DNA Clean and Concentrator-5 Kit, eluted using 21µL of elution buffer and stored at -20°C until library preparation.

Libraries were prepared exactly as described (Corces et al., 2017), using the adapter sequences listed in Suppl Table S9 (Buenrostro et al., 2013). After qPCR amplification, qPCR profiles were manually assessed to determine the additional number of cycles to amplify according to (Buenrostro et al., 2015). Libraries were purified using the Zymo DNA Clean and Concentrator-5 Kit and eluted in 11µL of H_2_O.

Concentrations and banding profiles were checked by Qubit and Tapestation. Sequencing (50bp paired-end) was conducted by Oxford Genomics to ≈15-20million reads -sufficient for open chromatin profiling of the chick genome.

ATACseq was also performed on pre-primitive-streak stage epiblast dissected from EGKXII-XIII embryos and on mature neural plate dissected from HH6-7 embryos (15-20 pieces of tissue per sample). Samples were collected in 1x Protease Inhibitor Cocktail in 1xPBS (cOmplete mini EDTA-free; Roche) and dissociated using a Dounce homogenizer. ATACseq was performed as described (Buenrostro et al., 2013; Buenrostro et al., 2015).

### Sample preparation for single cell RNA sequencing

Fertilized chicken eggs (Henry Stewart & Co. Ltd, Norfolk UK) were incubated at 38°C. The embryos were removed from the egg, staged, and pinned ventral side up on a resin plate in Tyrode’s saline. Embryos were dissected with a G-31 syringe needle. The endoderm and mesoderm were removed with a small volume of dispase (10mg/mL) before dissecting an ectodermal region spanning the neural plate and neural plate border (Figure 6A). Tissue samples were pooled from multiple embryos of the same stage prior to dissociation (HH4=∼55 embryos; HH6=∼45 embryos; HH8=10 embryos; HH9+=7 embryos). For cell dissociation, samples were incubated in FACSmax cell dissociation solution (Amsbio, T200100) with 30U/mL Papain (Sigma, P3125) at 37°C for 20 minutes. The tissue was pipetted 10 times every 5 minutes in order to facilitate mechanical dissociation. To stop enzymatic dissociation, an equal volume of resuspension solution (HBSS with 0.1% non-acetylated BSA (Invitrogen, 10743447), 1% HEPES, 1x non-essential amino acids (Thermo Fisher Scientific, 11140050) and 10μM rock inhibitor (StemCell Technologies, Y-27632) was added before passing cells through a 20μm filter (Miltenyi Biotech, 130-101-812). Samples were then centrifuged at 100rcf at 4°C for 5 minutes and resuspended in resuspension solution twice. Cells were then FAC sorted with 1uL 0.1mg/mL DAPI to remove dead cells and any remaining cell doublets. Immediately after FACS, samples were centrifuged and placed in 90% MeOH with 10% resuspension solution for storage.

Given the limited number of cells collected from individual dissection rounds and to obtain the required cell concentration for 10x, multiple rounds of dissection were carried out over successive days. Immediately prior to loading on the 10x Chromium controller, samples were pooled by stage and underwent three rounds of centrifugation at 1500rcf at 4°C for 10 minutes followed by rehydration with DPBS with 0.5% non-acetylated BSA and 0.5U/μl RNAse inhibitor (Roche 3335399001).

### Single cell library preparation and sequencing

Cells were loaded on the 10x Chromium controller with the aim of capturing 3000-5000 cells per stage. cDNA synthesis and library preparation were carried out using the Chromium Single-Cell 3′ Reagent v3 Kit (10× Genomics, Pleasanton, CA) according to the manufacturer’s protocol. This was carried out by the advanced sequencing facility at the Francis Crick Institute, London. Barcoded libraries were multiplexed and sequenced on the HiSeq 4000.

### RNAseq analysis

Raw sequencing data in FASTQ format underwent quality control analysis according to the pipeline published by (Blankenberg et al., 2010). Files were converted to Sanger format using FASTQ *Groomer*. The quality scores of reads were calculated using FASTQ *Summary Statistics*. Quality scores were measured as phred =-10 log_10_(p), where “p” is the estimated probability of a base being called incorrectly. Reads with a phred score of equal to or greater than 20 (i.e. 99% probability of a base being correctly identified) were kept, while poor quality reads with average phred scores of less than 20 were filtered out using FASTQ *Quality Filter*. The remaining paired-end reads were trimmed at the 3’ ends where the phred score drops below 20. All reads that passed quality control analysis were initially aligned to the chicken genome (assembly version Gallus_gallus-2.1, GenBank Assembly ID GCA_000002315.1) using TopHat (Trapnell, Pachter, & Salzberg, 2009). The raw data was re-analysed when the chicken genome was updated to assembly version gallus_Gallus-4.0, GenBank Assembly ID GCA_000002315.2 in 2013.

The node graft time course data together with previously published embryonic neural tissue RNAseq data (Trevers et al., 2018) were reanalysed using the *galGal5* genome assembly. Sequencing adapters and poor quality or N bases at 5’ and 3’ ends were trimmed from each sequence read using Trimmomatic-0.36 (Bolger, Lohse, & Usadel, 2014). Unpaired reads and those that were less than 36 bases long after trimming were also removed. The remaining reads were aligned to the *galGal5* genome using TopHat2 (Kim et al., 2013), alignment rates were 67.4% ± 12.9%. A custom GTF file was generated by adding additional gene ERNI (NM_001080874) annotation, taken from RefSeq, to Gallus_gallus-5.0 Ensembl release 94 GTF file. Transcripts were counted and normalized based on the custom edited GTF file using Cufflinks (Trapnell et al., 2012) program *cuffquant* and *cuffnorm* respectively. The output table contains transcript FPKM (Fragments Per Kilobase of transcript per Million mapped reads) values across 11 samples (node graft 4 time points and embryonic neural tissues with their controls). Sequence read tracks were computed using deepTools (Ramirez et al., 2016) *bamCoverage* (parameters *-bs 1 -- scaleFactor 10^6^/Library size --extendReads --samFlagInclude 66 –ignoreDuplicates --effectiveGenomeSize 1230258557*).

### Differential expression analysis

Differentially expressed transcripts were identified by comparing uninduced and induced tissue at each time point (5, 9 and 12h). Initially the “easyRNASeq” (Delhomme, Padioleau, Furlong, & Steinmetz, 2012), together with the Ensembl *galGal3* Gene Transfer File (GTF) were used to assemble and count transcripts from aligned reads. Differential expression analysis was then performed using 2 different methods: Cufflinks *cuffdiff* (Trapnell et al., 2012) and DESeq (Anders & Huber, 2010). The output from each analysis were merged; this identified 7745 transcripts that were differentially expressed with a log_2_ FC > 1.2 across all 3 time points (Suppl Table S15-16). Gene annotations were assigned to 4508 of these, corresponding to 2333 unique genes. Due to the incomplete nature of the *galGal3* chicken genome, 3237 transcripts were left unannotated.

This process was repeated using the Ensembl and UCSC *galGal4* GTF when they were released. For the Ensembl constructed transcripts, gene annotations and chromosome locations were added using Ensembl Biomart data. For UCSC constructed transcripts, gene annotations and chromosome locations were added using UCSC *galGal4* GTF and annotation data from the UCSC table browser. Fully annotated transcripts from either Ensembl or UCSC lists were combined to provide the most comprehensive set of annotations. This reanalysis identified 8673 differentially expressed transcripts across the 3 time points (Suppl Table S17-18). Gene annotations were assigned to 7184 of these, corresponding to 4145 unique genes, but 989 transcripts still remained unannotated.

The expression data (FPKM table) generated based on *galGal5* genome assembly was analysed in R-3.5.1 environment. FC was calculated from the levels of expression in the induced relative to the corresponding uninduced tissue, or neural relative to non-neural tissue samples. Genes with FPKM >10 in an induced or neural tissue and FC >1.5 are defined as upregulated, whereas genes with FPKM >10 in an un-induced/non-neural tissue and FC <0.5 are defined as downregulated. For the time point 0, a gene is defined as upregulated when 0h uninduced FPKM >10, FC >1.5 against 5h induced tissue, and 5 hour uninduced FPKM value is over 10 and bigger than the value in 5h induced tissue. The FPKM expression levels of the candidate GRN members are provided in Suppl Table S2.

### NanoString data analysis

Raw NanoString data were analysed in Microsoft Excel according to NanoString guidelines. Data were checked to ensure 600 fields of view were counted and that binding density values fell within the 0.05-2.25 range. Next, the geometric mean (geomean) of 6 positive control probes was calculated for each assay and averaged across the entire data set. From these, a positive lane normalization factor (PLNF) was calculated by dividing the average geomean by the geomean for each assay. Then, negative and endogenous probe counts were multiplied by their respective PLNF to normalize for differences in numbers of reporter and capture probes between assays. Next, the geomean of housekeeping (HK) genes ACTB and GAPDH was calculated for each assay and averaged across all assays. The average HK geomean was then divided by the HK geomean for each lane, to generate a lane specific normalization factor (LSNF). Then, negative and endogenous probe counts were multiplied by the LSNF to normalize for cell number differences between assays. For each assay, the mean plus two times standard deviation was calculated from the 8 normalized negative control probes. This value was subtracted from all normalized endogenous probe counts to remove background noise levels. Any transcript counts that became negative as a result, were reset to zero. Next, the normalized expression counts were set to 1 when the counts are equal to zero. Average counts for each probe and standard deviation were calculated across triplicate assays. Finally, FC for each probe was calculated by dividing the induced average by uninduced average. Differential expression was defined as FC of >1.2 (upregulation) or <0.8 (downregulation). The statistical significance was calculated using a Two-tailed Type 2 T-Test with a P-value of 0.05. The result is provided in Suppl Table S3.

### ChIPseq analysis

Single-end sequence reads were quality checked and trimmed using Trimmomatic-0.36 with the same criteria as RNAseq analysis and aligned to the *galGal5* genome using bowtie2 (Langmead & Salzberg, 2012) with default parameters. The alignment rates were 92.3% ± 6.5%. PCR duplicates were removed using Piccard *MarkDuplicates* (parameter *REMOVE_DUPLICATES=TRUE*). Peak calling was computed using MACS2 (Zhang et al., 2008) *callpeak* (parameters *-f SAM -B --nomodel --broad --SPMR -g 1230258557 -q 0.01*) with genomic input as control. Signal tracks were computed using deepTools *bamCompare* (parameters *--scaleFactorsMethod SES --operation log2 -bs 1 --ignoreDuplicates*) and displayed using R package trackViewer (Ou & Zhu, 2019). CTCF ChIPseq data (Kadota et al., 2017) were processed with the same pipeline described here. The resulting file (bed format) from peak calling was used as an input file to the pipeline for GRN.

### ATACseq analysis

Paired-end sequence reads were quality checked and trimmed using Trimmomatic-0.36 with the same criteria as RNAseq and aligned to the *galGal5* genome using bowtie2 (parameters *-X 2000 --sensitive-local*). The alignment rates were 92.36% ± 5.33%. PCR duplicates were removed using Piccard *MarkDuplicates* (parameter *REMOVE_DUPLICATES=TRUE*). Coverage tracks were generated using deepTools bamCoverage (parameters *-bs 1 --scaleFactor 10^6^/library size --extendReads --samFlagInclude 66 --ignoreDuplicates --effectiveGenomeSize 1230258557*).

### Single cell RNAseq data analysis

scRNAseq alignment and downstream analysis was run using Nextflow (version 20.07.1) (Di Tommaso et al., 2017) and Docker for reproducibility. All of the required packages and respective versions are found within the Docker container alexthiery/10x_neural_tube:v1.1. The full analysis pipeline, including documentation, can be found at https://github.com/alexthiery/10x_neural_tube.

In order to obtain the recommended 50k reads/cell, libraries were sequenced twice, with reads from both flow cells pooled during sequence alignment. scRNAseq reads were demultiplexed, aligned and filtered using Cell Ranger (version 3.0.2, 10x Genomics). Reads were aligned to *galGal5*, using a custom edited GTF annotation file as described in “RNAseq analysis” section. Prior to alignment, MT, W and X chromosome genes in the g*alGal5* GTF file were prefixed accordingly for simple identification downstream. Coordinates for W chromosome genes in *galGal6* were also transferred to *galGal5* and prefixed in the GTF.

Quality control, filtering, dimensionality reduction and clustering were carried out in Seurat (version 3.1.5) (Stuart et al., 2019). We first applied a modest filtering threshold, removing cells with greater than 15% UMI counts from mitochondrial genes and cells with fewer than 1k or greater than 6k unique genes. After filtering 8652 cells remain in the dataset (HH4=2745; HH6=1823; HH8=1750; HH9+=2334). Throughout subsequent clustering steps, we measure total UMI counts, total gene counts and percentage mitochondrial content for each cluster and remove any clear outlier clusters. Following initial clustering, cell clustering was dominated by the expression of a few W chromosome genes and we therefore removed W chromosome genes. We then identified and removed contaminating cell clusters, including putative mesoderm and primordial germ cells. Percentage mitochondrial content, cell cycle and sex were all regressed out during scaling of the final dataset.

PCA was used as an initial dimensionality reduction step, with the top 15 principal components used for clustering. We then calculated k-nearest neighbours in order to embed a shared nearest neighbour (SNN) graph. The Louvain algorithm was used to partition the SNN graph according to the Seurat default parameters. At each clustering step, we visualized multiple resolutions using the R CRAN clustree package (Zappia & Oshlack, 2018) and determined the optimal resolution manually based on cluster stability.

We unbiasedly identified modules of co-correlated genes using the Antler gene modules algorithm (https://github.com/juliendelile/Antler) (Delile et al., 2019). For this we kept genes which have a Spearman correlation greater than 0.3 with at least 3 other genes. Ward’s hierarchical clustering is then carried out on the Spearman gene-gene dissimilarity matrix of the remaining genes. Iterative clustering by the Antler algorithm selects modules based on their consistency. In order to identify gene modules that explain the greatest amount of variation between our cell clusters, we filtered modules for which fewer than 50% of genes are differentially expressed (logFC > 0.25, adjusted p-value < 0.001) between at least one cell cluster and to the remaining dataset. Differential expression tests were carried out using Seurat FindAllMarkers function (Wilcoxon test).

Neural cell clusters were subset from the parent dataset using a candidate marker approach. PC1 inversely correlates with developmental stage, therefore cells were subsequently ranked according to their position along PC1. GRN components were grouped based on the timing of their initial expression following neural induction. Housekeeping genes (*GAPDH*; *SDHA*; *HPRT1*; *HBS1L*; *TUBB1*; and *YWHAZ*) were used as a control. A general additive model was used to estimate the changes in scaled expression across the ranking of PC1, with a smoothing spline fitted by REML.

### Pipeline for GRN

Two hundred and thirteen transcription regulators were selected from the RNAseq time course data (0, 5, 9 and 12h). The data from NanoString nCounter assay provided fine time course expressions (1, 3, 5, 7, 9, 12h after node graft) of the candidate components of the GRN. It not only is a supportive dataset to the RNAseq expression data for the time points 5, 9 and 12h, but also provides gene expression levels for 1, 3, 7h for the GRN. The details for differential expression analyses and thresholds are described in the “Differential expression analysis” sections. The integrated time course gene expression profile can be found in Suppl Table S1. There were 180 transcription regulators that were found differentially expressed in both RNAseq and NanoString datasets and thus remained in the list of the candidate GRN components.

To define the neural induction regulatory loci, the genomic coordinates of the GRN candidate members were extracted from Gallus_gallus-5.0 Ensembl release 94 GTF file. The in-house script searches upstream and downstream of the candidate members up to 500kb for the peaks with highest *signalValue* from CTCF ChIPseq peak calling result file. The coordinates of regulatory loci are listed in Suppl Table S4.

Different types/indices of putative regulatory sites (Figure S2A) were obtained from the ChIPseq H3K27 peak calling result files using BedTools (Quinlan & Hall, 2010). The command *intersect* reports the overlapping peaks between the ChIPseq results, while *intersect* with parameter *-v* reports the non-overlapping peaks between a set of ChIPseq outputs. The command *merge* (parameters *-d 100 -c 1,4 -o count,collapse*) was used to merge and generate a unique peak set from a subset of results. “Activation” regulatory sites, as an example, were merged from indices 1-3. The coordinates of the regulatory loci in Suppl Table S4 were then used as an input to the in-house script to extract the coordinates of regulatory sites of the GRN candidate members. FASTA format sequences of these regulatory sites were also extracted from *galGal5* genome assembly Ensembl release 94 for the transcription factor binding site screening.

A transcription factor binding motif library of the GRN candidate members was extracted from JASPAR2020 CORE non-redundant database (Fornes et al., 2020). SALL1 and GRHL3 consensus binding motifs were obtained from (Karantzali et al., 2011; Klein et al., 2017) and converted into meme motif format. A total of 91 transcription factor binding motifs were curated as an input library for running binding motif screening on the putative regulatory sites (FASTA format files) using MEME suite FIMO (Grant, Bailey, & Noble, 2011). The in-house script selects predicted biding sites from FIMO outputs with confidence score >10 and calculates the genomic coordinates of the binding sites. The resulting outputs are provided in Suppl Table S5-7.

The in-house script integrates gene expression profile and transcription factor binding profile within the putative regulatory sites that undergoing activation (indices 1-3 and 7) to generate the neural induction GRN. A positive regulatory interaction is modelled when the regulator and the target gene are both up or down regulated at one time point, whereas a negative regulation is predicted when the regulator and the target gene have opposing expression profiles (Figure S3A-B). Duplicated interactions derived from the bindings of a regulator to the multiple regulatory sites of a target gene were removed. The putative regulatory sites from 5h ChIPseq data were used for generating the GRN at time point 1, 3 and 5h. The sites from 9h ChIPseq were used for the GRN at time point 7 and 9h. The in-house script generates files, including Model Hierarchy CSV and Time Expression XML Data, for visualizing the GRN using BioTapestry (Longabaugh et al., 2005, 2009; Paquette et al., 2016). The BED files were also generated for uploading to the UCSC browser for visualizing the GRN related data. Five genes, including BAZ1A, GATAD2B, CRIP2, PBX4 and SALL3, were excluded from the GRN, because they have neither input nor output interactions with other members of GRN. This gives the final GRN with 175 components in the network.

DREiVe (Sosinsky et al., 2007) was used to search for conserved elements within 500kb up- and down-stream of the 175 GRN components. The default settings were used to compare between 6 species: human (*hg38*), mouse (*mm10*), rat (*rn6*), golden eagle (*aquChr2*), zebrafish (*danRer10*) and chicken (*galGal5*). Elements that are conserved between a minimum of 4 species were selected and displayed on the UCSC browser.

To generate a subnetwork of a selected target gene BRD8, the transcription factors, predicted to bind to the regulatory sites that undergoing either activation or repression (indices 1-7), were extracted from the FIMO outputs. BedTools *merge* (parameters *-d 500 -c 1,4 -o count,collapse*) was used to check the overlaps of the regulatory sites across 3 time points (5, 9 and 12h). Each unique regulatory site was treated as one independent component in the network. The in-house script generates the same output files as the main GRN for visualizing using BioTapestry.

## Data and Code Availability

Raw and processed data generated during this study are available either as supplementary files or have been deposited to ArrayExpress. The accession number for the RNAseq data reported in this paper is ArrayExpress: E-MTAB-10409. The accession number for the ChIPseq data reported in this paper is ArrayExpress: E-MTAB-10424. The accession number for the ATACseq data reported in this paper is ArrayExpress: E-MTAB-10426 and E-MTAB-10420. The accession number for the single cell RNAseq data reported in this paper is ArrayExpress: E-MTAB-10408.

Analysis scripts are available at https://github.com/grace-hc-lu/NI_network and https://github.com/alexthiery/10x_neural_tube

## Supporting information

Main text, figures and methods

## Acknowledgements

This study was funded by grants from NIH (R01 MH 60156), MRC (G0400559), Wellcome Trust (063988) and BBSRC (BB/R003432/1 and BB/K007742/1) to CDS and BBSRC (BB/K006207/1) to AS. The work of NML is supported by the Francis Crick Institute which receives its core funding from Cancer Research UK (FC010110), the UK Medical Research Council (FC010110), and the Wellcome Trust (FC010110). The single-cell RNAseq data analyses were performed using infrastructure from the Crick Scientific Computing science technology platform. NML is a Winton Group Leader in recognition of the Winton Charitable Foundation’s support towards the establishment of the Francis Crick Institute.

